# Unsupervised Representation Learning Reveals Individualized Neurophysiological Profiles

**DOI:** 10.64898/2026.02.10.705127

**Authors:** Maxence Lapatrie, Jason da Silva Castanheira, Idil Aydin, Sylvain Baillet

**Affiliations:** Dept. of Electrical and Computer Engineering, McGill University, Montreal, Canada; McConnell Brain Imaging Centre, Montreal Neurological Institute, Montreal, Canada; Div. of Psychology and Language Sciences, Inst. for Cognitive Neuroscience, University College London, London, United Kingdom; Research Centre, University of Montreal Health Centre (CRCHUM), Montreal, Canada; Dept. of Neuroscience & Medicine, University of Montreal, Montreal, Canada

**Keywords:** human neurophysiology, magnetoencephalography, brain fingerprinting, neurophysiological profiling, autoencoders

## Abstract

Human brain activity contains stable, individual-specific features that persist over months to years, forming neurophysiological profiles. Most model-based profiling approaches use participant labels or supervised objectives, making it difficult to determine whether successful differentiation reflects stable biology or exploitable idiosyncrasies. We introduce a participant-agnostic autoencoder framework that derives profiles from brief resting-state magnetoencephalography (MEG) segments using reconstruction as sole training objective. Discriminative profiles emerged from the learned latent space without participant labels. Within-session, autoencoder profiles reached 93.3% accuracy at 120 s, exceeding functional-connectivity, spectral, and contrastive baselines with recordings as short as 14 s when participant-specific anatomy was withheld from source reconstruction. Differentiation generalized above chance across recording sessions (between-session accuracy 49.5% for the pretrained autoencoder). Profiles also predicted age more accurately than baselines (*r*^2^=0.318), and the decoder enabled perturbation-based sensitivity analyses in spectral and connectivity spaces. This establishes participant-agnostic representation learning as a scalable and interpretable profiling.

## Introduction

Resting neurophysiological activity contains stable, individual-specific signatures that can persist over months to years and support participant differentiation from brain recordings alone, often referred to as brain fingerprinting. Across modalities, including fMRI and magneto-/electroencephalography (M/EEG), individual differentiation has been demonstrated using functional connectivity, network topography and spectral structure^1–3^. Short-term stability over seconds is expected from temporal autocorrelation^4–6^, but stability across days to months suggests that enduring individual traits are embedded in ongoing brain dynamics rather than arising solely from transient brain states.

Because M/EEG resolves neural activity at millisecond timescales, it provides access to spatiotemporal structure that is not captured by spatial connectivity alone. Participant differentiation has therefore been pursued using connectivity, spectral content and temporal organization^7–9^. The relevance of such profiles extends beyond identification: features derived from functional and spectral representations can be heritable^10,11^ and have been associated with neurodegenerative conditions, including Parkinson’s and Alzheimer’s disease^12–14^.

Despite this promise, most profiling methods rely on model-free representations such as functional connectomes or power spectra. These representations are interpretable and physiologically grounded, but they impose predefined transforms that compress the signal and constrain the feature space a priori. Model-based approaches can instead learn mappings from time series or predefined features into latent spaces from which profiles are extracted^15–24^. In principle, deep neural networks can capture non-linear spatiotemporal dependencies beyond standard spectral or connectivity analyses and may support generalization across datasets and acquisition protocols.

Many existing model-based profiling methods, however, optimize participant separability directly, using classification heads or contrastive objectives that draw embeddings from the same participant together and push embeddings from different participants apart. These objectives can be effective for differentiation, but they shape the geometry of the latent space around decision boundaries rather than around the data-generating structure. They can therefore encourage shortcut learning, whereby models exploit anatomical differences, residual environmental or physiological artifacts after preprocessing and quality control, or session-specific recording features instead of stable neurophysiological traits^25–29^. In MEG specifically, such shortcuts could exploit participant-specific head geometry source leakage, within-session head position, or site- and session-specific artifacts that survive preprocessing. None of these reflect stable neurophysiological traits. High within-dataset accuracy may therefore not guarantee robustness across sessions, devices, preprocessing pipelines or cohorts. This motivates the surrogate-anatomy and between-session tests reported below.

A profiling framework intended for general use should instead learn representations of ongoing brain activity such that participant differentiation emerges as a consequence, rather than as the training target. Unsupervised learning provides a natural route, but remains underexplored for neurophysiological profiling. Sparse dictionary learning has improved individual differentiation in fMRI connectomes without participant labels, but still operates on predefined feature spaces that are already discriminative^17,21^.

Here, we propose an unsupervised, participant-agnostic autoencoder framework to derive neurophysiological profiles directly from MEG source time series. The same strategy is, in principle, extensible to other brain-recording modalities such as EEG and potentially fMRI. Autoencoders compress high-dimensional data into latent representations while preserving information required for reconstruction. We hypothesized that reconstructing short segments of ongoing activity would require the model to encode stable structure that recurs across segments, whereas transient state-dependent variation would be attenuated by temporal averaging in the latent space.

We trained an autoencoder to encode and decode brief segments of resting-state brain activity without participant identities or supervised differentiation objectives (Figure 1). Although the model was optimized only for reconstruction, stable and highly discriminative individual structure emerged in the learned latent space. We formed neurophysiological profiles by temporally pooling latent representations across short windows, suppressing window-specific variability while retaining consistent individual structure.

**Figure 1:**
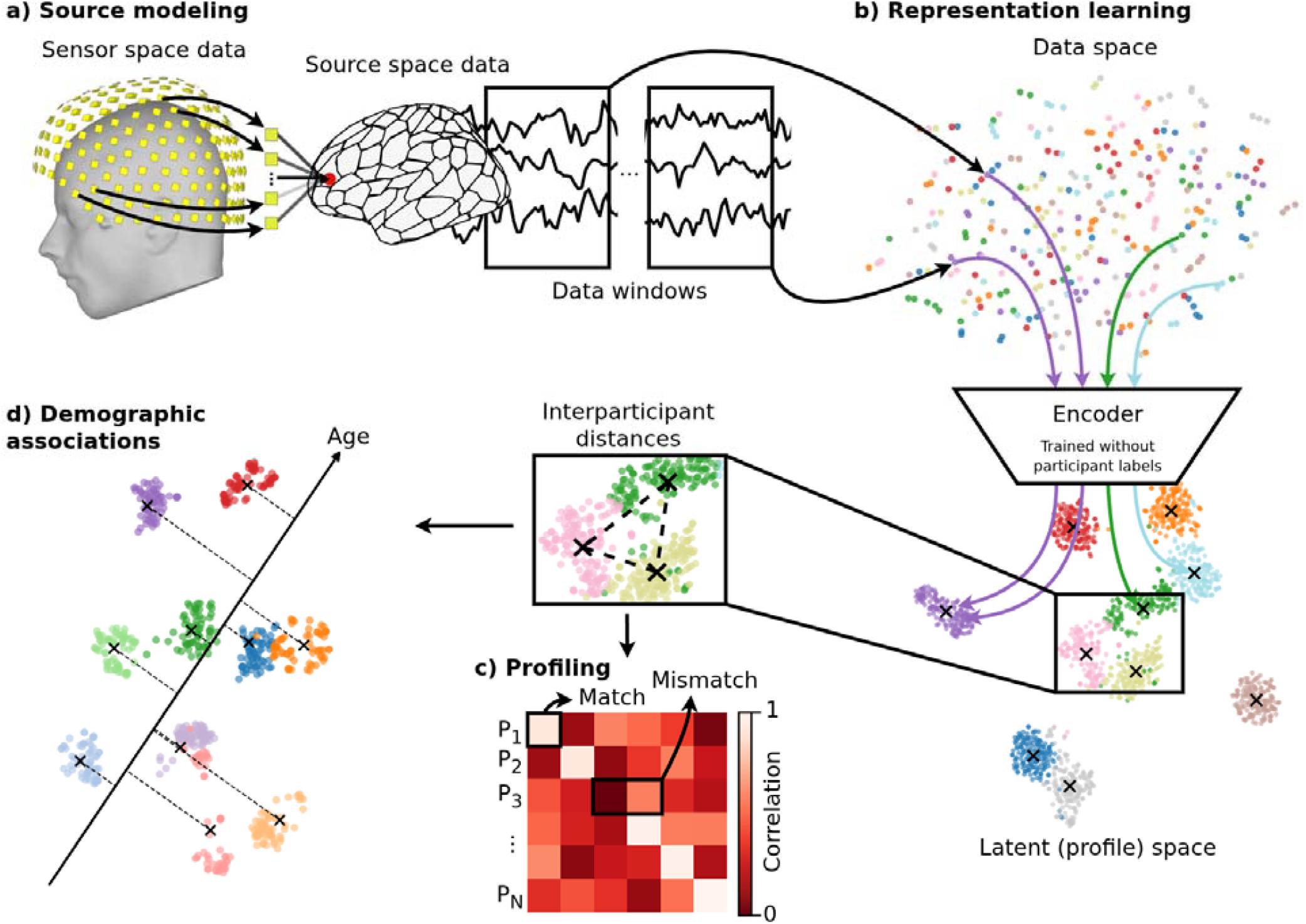
Conceptual overview of participant-agnostic neurophysiological profiling. a) Source modeling. Resting-state MEG sensor data were projected to source space, parcellated into cortical regions of interest (ROIs), and segmented into 6-s windows. Source reconstruction was performed in a common anatomical template space using either participant-specific (“native anatomy”) or shared (“surrogate anatomy”) anatomical information. b) Representation learning. Window-level source signals were mapped to latent (profile) space by an autoencoder trained without participant labels. Latent representations from multiple windows were temporally pooled to form participant-level neurophysiological profiles. Colors indicate participant labels for visualization only. c) Profiling. Participant differentiation was performed by computing pairwise Pearson correlations between profiles derived from independent data segments. A correct match occurs when the highest correlation for a participant falls on the diagonal of the similarity matrix. d) Demographic associations. Low-dimensional visualization of the latent profile space (visualization only). Linear regression directions in profile space were used to predict demographic variables such as age. Participant centroids are projected onto the regression axis to illustrate predicted values.

Across independent MEG datasets, these profiles were the most discriminative of the tested approaches for within-session differentiation under surrogate anatomy, outperforming baselines from recordings as short as 14 s and remaining robust when participant-specific anatomy was withheld from source reconstruction. For between-session differentiation, the profiles generalized above chance and remained competitive with model-free baselines while avoiding the collapse exhibited by the contrastive encoder. Because the model includes a generative decoder, latent perturbations can be projected back into interpretable spectral and functional-connectivity features, addressing a central limitation of many model-based approaches. The framework thus establishes participant-agnostic unsupervised learning as a scalable and interpretable approach to neurophysiological profiling that prioritizes signal structure over task-or dataset-specific separability.

## Results

Unless specified otherwise, results are reported for resting-state MEG data from the Cambridge Centre for Ageing and Neuroscience (Cam-CAN) dataset^30^. Data were processed in two source spaces: native anatomy, in which participant-specific anatomy informed source reconstruction, and surrogate anatomy, in which a single held-out anatomy was shared across participants to assess robustness to anatomical information.

We trained an autoencoder (AE) and a contrastive encoder (CONT), and subsequently fine-tuned the AE using the reconstruction objective (fine-tuned AE). All models were trained in the native anatomy source space and evaluated on a single independent test set of 100 participants, reconstructed twice—once with participant specific (native) anatomy and once with a shared surrogate anatomy. No participant identities entered training. The autoencoder never receives identity information regardless of the subset, and the contrastive encoder used identities only from the training set, never from the test set. Unless otherwise noted, accuracies are reported as mean ± SD over 100 repetitions of 90% participant subsampling without replacement (Methods).

### Autoencoder Output Validation

Before using the learned latent representations for profiling, we asked whether the autoencoder preserved essential properties of the underlying brain recordings. Because the model was trained only to reconstruct short time-series windows, its relevance depends on whether reconstruction fidelity extends beyond raw signal similarity to established neurophysiological features. We therefore evaluated preservation of temporal structure, spectral content and functional connectivity, and identified frequency ranges or features that were less reliably recovered.

We encoded and reconstructed 10 non-overlapping 6-s windows per participant from an independent test set (N = 100), and compared original and reconstructed signals across multiple feature-level metrics.

The decoder reconstructed input windows with substantial fidelity (Figure 2). Across channels, time-series reconstruction fidelity was r = 0.73 ± 0.02, with lower performance in low-amplitude channels. AEC matrices computed from reconstructed windows closely matched those from the original signals (broadband 0.5-45 Hz: r = 0.70 ± 0.03). AEC fidelity was high across all frequency bands except gamma (30-45 Hz), where fidelity decreased by Δr = 0.21 relative to broadband (Extended Data Figures 1-2).

**Figure 2:**
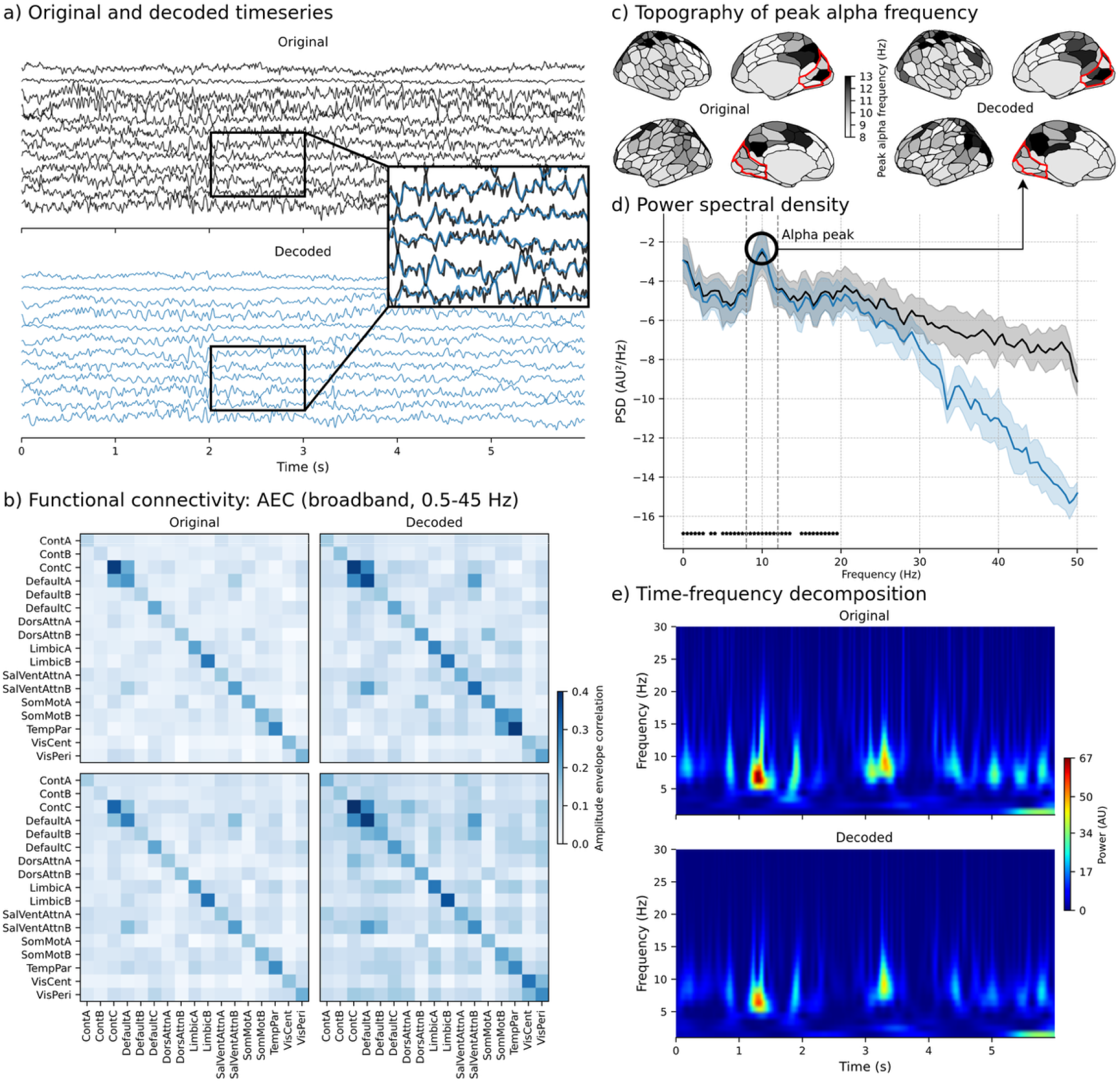
Generative validation of autoencoder reconstructions. a) Original and decoded time series. Example 6-s source-level window from visual peripheral ROIs (top: original; bottom: decoded). Signals are z-scored per window using the global mean and standard deviation to preserve relative topography. Overall temporal structure is retained, with attenuation of high-frequency detail. b) Functional connectivity. Amplitude-envelope correlation (AEC; broadband 0.5-45 Hz) computed on the same 6-s window for original and decoded data, averaged across Yeo-17 networks. Participant-specific connectivity structure is preserved. c) Topography of peak alpha frequency. Cortical maps of peak alpha frequency (PAF) averaged over 10 consecutive windows (60 s) from one participant (left: original; right: decoded), showing close spatial correspondence. Visual peripheral ROIs used in d are outlined in red. d) Power spectral density. Mean ± SD PSD across visual peripheral ROIs for original (black) and decoded (blue) signals. The alpha peak (8-13 Hz) is preserved. Frequency bins marked with stars have substantial evidence for spectral similarity (*BF*_01_ ≥ 3), supporting the null hypothesis of no difference between original and reconstructed signals. e) Time-frequency decomposition. Morlet wavelet time-frequency representations (1-30 Hz; 3 cycles) averaged over somatomotor ROIs for the same 6-s window (top: original; bottom: decoded), illustrating preservation of transient spectral bursts. This panel is shown for qualitative comparison.

Power spectral density (PSD) estimates within Yeo-17 networks^31,32^ showed substantial evidence for similarity (*BF* _01_≥ 3 ) across frequencies below approximately 25 Hz, although the preserved bandwidth varied across ROIs (Extended Data Figures 3-4). Peak alpha frequency (PAF) topographies were moderately preserved (r = 0.54 ± 0.15), and time-frequency structure was retained, with slight attenuation of high-frequency power in reconstructed windows (Extended Data Figures 5-6).

### Profiling

We next compared model-free and model-based approaches to neurophysiological profiling. Model-free profiles^7–9^ were derived from either functional connectivity (AEC) or spectral information (PSD estimates). Model-based profiles were obtained by encoding fixed-length windows with the AE, fine-tuned AE or CONT and averaging consecutive latent representations within each participant.

#### Effect of Segment Length on Differentiation Accuracy

If stable individual-specific structure is encoded in the latent space, differentiation accuracy should improve as longer segments are used to average out transient state-dependent variability. A practical goal of neurophysiological profiling is also to minimize the amount of clean data required for reliable differentiation. We therefore quantified how differentiation accuracy scales with recording duration across model-free and model-based approaches, using segment lengths from 10 to 120 s.

In the surrogate anatomy condition, accuracy increased monotonically with segment length for all approaches (Figure 3a). At 120 s, the AE and fine-tuned AE achieved the highest accuracies (93.0 ± 0.9% and 93.4 ± 1.0%, respectively), exceeding functional connectivity (87.0 ± 1.1%), PSD (78.3 ± 1.6%) and CONT (78.6 ± 1.6%).

**Figure 3:**
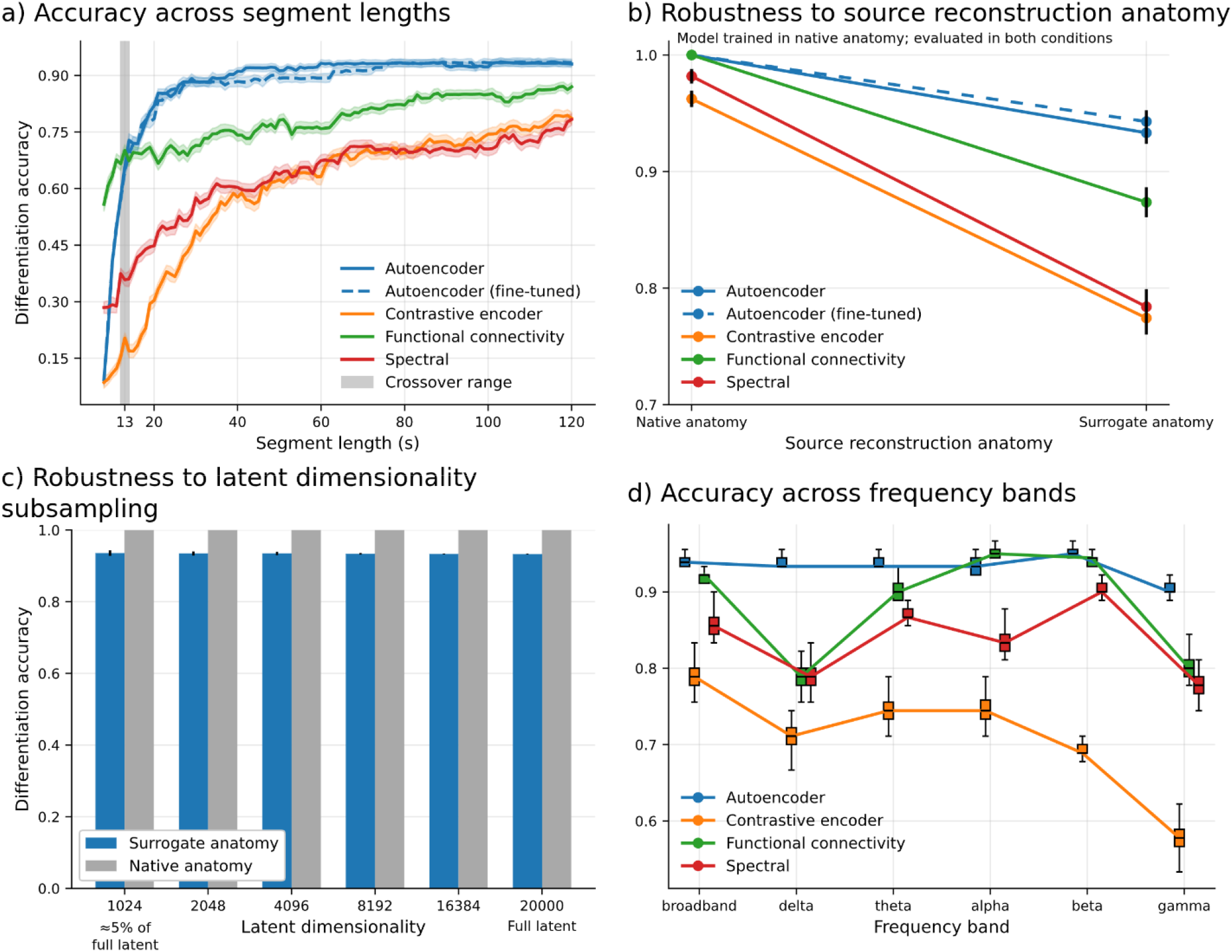
Within-session participant differentiation in Cam-CAN. a) Accuracy across segment length. Top-1 participant differentiation accuracy as a function of recording duration (10-120 s), computed by maximal Pearson correlation between profiles derived from the beginning and end of each recording (N = 100). Chance level is 1%. Curves show mean accuracy; shaded bands indicate bootstrap SD over 100 resamples of 90% of participants. The shaded vertical region highlights the empirical crossover range (12-14 s) beyond which autoencoder-based profiles consistently outperformed baseline methods. b) Robustness to source reconstruction anatomy. Differentiation accuracy at 120 s when models trained in native anatomy space were evaluated using either participant-specific (native) or shared (surrogate) anatomy during source reconstruction. Autoencoder-based profiles exhibit smaller accuracy reductions than model-free and contrastive baselines. Error bars denote bootstrap SD. AE, fine-tuned AE and FC showed no variance across bootstrap subsets in the native condition, resulting in non-visible error bars. c) Robustness to latent dimensionality subsampling. Differentiation accuracy after randomly subsampling the autoencoder latent space to reduced dimensionalities (100 draws per dimensionality). Accuracy remains stable down to 1,024 dimensions (∼5% of the full latent space). d) Accuracy across frequency bands. Differentiation accuracy at 120 s after band-pass filtering inputs into canonical frequency bands. Model-free approaches show frequency-specific performance, whereas autoencoder-based profiles remain relatively invariant across bands. Error bars denote bootstrap SD. Results are shown for surrogate anatomy.

At 10 s, differentiation accuracy remained above chance for all methods, with FC performing best (63.3 ± 1.7%), followed by AE (40.5 ± 1.7%), PSD (29.2 ± 2.1%) and CONT (11.0 ± 1.3%). By 30 s, AE performance largely plateaued (88.3 ± 1.3%), exceeding FC by 16.8 percentage points (adjusted p < 0.001). AE outperformed all other approaches for segments of 14 s and longer (adjusted p < 0.001).

#### Robustness to Anatomy Alignment

A central concern in neurophysiological profiling is the extent to which individual differentiation relies on participant-specific anatomy rather than functional dynamics. Because source reconstruction incorporates anatomical information, strong differentiation performance could in principle reflect anatomical idiosyncrasies rather than neurophysiological traits. We therefore evaluated robustness when participant-specific anatomy was withheld from MEG source mapping by applying a shared surrogate anatomy to all participants.

In the *native anatomy* space, with 120-s recordings, accuracies reached ceiling levels for FC, AE, and fine-tuned AE (100.0 ± 0.0%), with slightly lower performance for PSD (98.2 ± 0.6%) and CONT (96.2 ± 0.7%). When evaluated in *surrogate anatomy* space, accuracies decreased to 87.4 ± 1.3% (FC), 78.4 ± 1.5% (PSD), 93.3 ± 0.9% (AE), 94.3 ± 1.0% (fine-tuned AE), and 77.4 ± 1.4% (CONT). These 120-s surrogate-anatomy accuracies differ slightly from those reported in the segment-length analysis above because the two analyses are independent experiments that drew separate bootstrap resamples under different random seeds. The underlying test and data are identical.

Autoencoder-based profiles showed the smallest performance drops across anatomical conditions (Figure 3b), with reductions of 6.7 percentage points (AE) and 5.7 percentage points (fine-tuned AE), compared with 12.6-19.8 percentage points for other approaches. All methods remained well above chance.

#### Effect of Post Hoc Latent Dimensionality Reduction

Although the autoencoder produces high-dimensional latent representations, stable participant-specific information may be concentrated in a lower-dimensional subspace. To assess redundancy, we randomly subsampled *d* ∈ {1024,2048,4096,8192,16384} dimensions from the 20,000-dimensional latent profiles and recomputed differentiation accuracy without retraining. Profiles were derived from 120-s recordings.

Differentiation accuracy remained stable across tested dimensionalities (Figure 3c). In the native anatomy space, AE performance remained at 100.0 ± 0.0%. In the surrogate anatomy space, accuracy was 93.3 ± 0.1% at full dimensionality and 93.6 ± 0.8% at 1,024 dimensions, corresponding to approximately 5% of the original latent space.

#### Frequency Band Contributions

Model-free profiling methods typically show strong frequency-specific effects, particularly in the alpha-beta range^8,9^ (alpha, 8-13 Hz; beta, 13-30 Hz). To determine whether autoencoder-based profiles rely preferentially on specific frequency bands or integrate information more broadly, we evaluated differentiation accuracy after band-pass filtering input signals into six canonical frequency bands, without retraining the models. Models were evaluated in the surrogate anatomy condition with 120-s recordings.

Model-free approaches peaked in the theta-beta range (theta, 4-8 Hz; alpha, 8-13 Hz; beta, 13-30 Hz), as expected (Figure 3d). FC achieved its highest accuracy in the alpha band (95.3 ± 1.1%), and PSD peaked in beta (90.3 ± 1.0%). CONT performed best in broadband (79.0 ± 1.6%) and showed reduced accuracy in isolated bands. In contrast, AE accuracy was largely invariant across frequency bands, remaining within 5 percentage points across bands.

#### Generalizability to Between-Session Profiling

Robust neurophysiological profiles should generalize across recording sessions separated by days to months, despite changes in brain state, environment and recording conditions. To test whether representations learned from within-session data capture enduring individual traits, we evaluated between-session differentiation in an independent longitudinal dataset.

Using the OMEGA dataset^33^, we trained models on 200 single-session participants and evaluated performance on 78 participants with multiple sessions (Extended Data Figure 7). We tested both within- and between-session profiling with 120-s recordings from which we derived the profiles. Within-session profiling yielded 100.0 ± 0.0% accuracy for all methods. Between-session accuracy decreased for all approaches, to 53.7 ± 2.1% (FC), 53.2 ± 1.9% (PSD), 49.5 ± 2.0% (AE), 56.2 ± 2.3% (fine-tuned AE), with the contrastive encoder falling to 13.1 ± 1.4%. The four non-contrastive methods were closely grouped, with comparable drops of approximately 45-50 percentage points and no method separating clearly from the others. The lead of the fine-tuned AE reflects its status as an upper bound of the autoencoder approach to profiling rather than genuine between-session superiority. The contrastive encoder, by contrast, showed a substantially larger decrease (86.9 percentage points). All methods remained above chance. The salient between-session finding is therefore that reconstruction-based profiling generalizes above chance and avoids the collapse exhibited by the participant-supervised contrastive objective.

### Interpretability

#### Latent Space Stability Analysis

Interpretable learned representations should exhibit stable, participant-specific structure. Because the autoencoder latent space is unordered, we first asked whether latent components could be ranked by stability across participants and used to guide systematic perturbation analyses.

Principal Component Analysis (PCA) revealed a clear stability hierarchy in the latent space. Intraclass correlation coefficients decreased monotonically with component index, following a power-law relationship (r^2^ = 0.747), with leading components reaching ICC(3, 1) ≈ 0.95 (Methods). Occlusion of individual components followed by decode-reencode cycles indicated that leading components define consistent, non-scrambled directions in latent space, supporting their use for perturbation-based analyses (Extended Data Figure 8b).

#### Feature Sensitivity Derived from Decoder-based Perturbations

A frequent critique of model-based approaches is limited interpretability, because latent dimensions do not map explicitly onto neurophysiological features. Here, the generative decoder enabled direct perturbation-based assessment of how latent structure propagates to interpretable signal features.

We reconstructed 5,226 latent vectors from test participants after setting the first principal component (PC1) to zero and quantified feature sensitivity as the mean absolute z-score deviation relative to participant-specific baseline distributions. Zeroing PC1 induced structured and reproducible changes in both functional connectivity and spectral features. For FC, the largest band-wise effects occurred in the delta band, with network-level sensitivity concentrated in dorsal attention networks. For PSD, sensitivity was more broadly distributed, with strongest effects in beta and broadband activity (Figure 4).

**Figure 4.**
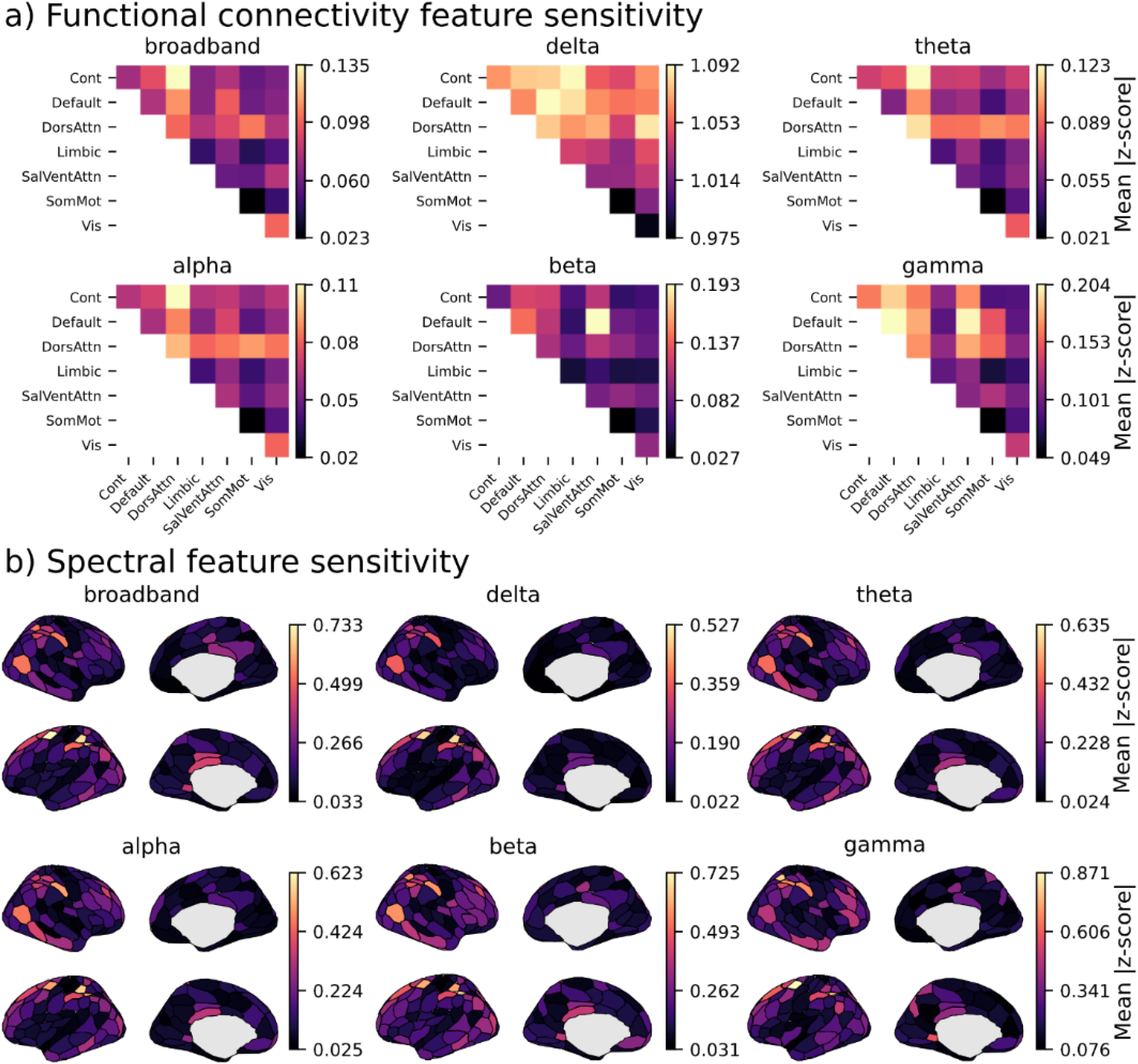
Decoder-based sensitivity of latent components to interpretable neurophysiological features.a) Functional connectivity feature sensitivity. Mean absolute z-score change in amplitude-envelope correlation after zeroing PC1, summarized across Yeo-17 networks and canonical frequency bands. b) Spectral feature sensitivity. Mean absolute z-score change in PSD after zeroing PC1, projected to cortical parcels and shown by frequency band. Warmer colors indicate greater sensitivity; values are normalized relative to participant-specific baseline variability.

### Demographic Associations

If learned profiles capture biologically meaningful individual variability, they should encode information relevant to demographic factors known to influence brain activity. We therefore evaluated whether autoencoder-based profiles predict age and sex, and whether latent-space geometry constrains linear demographic prediction. Performance was compared with model-free and participant-supervised baselines.

Latent-space geometry of the autoencoder was assessed by comparing geodesic distances, estimated as shortest-path distances on a k-nearest-neighbor graph, with Euclidean distances. Euclidean distances approximated geodesic distances substantially better in the surrogate anatomy space (r^2^ = 0.914 ± 0.003) than in the native anatomy space (r^2^ = 0.849 ± 0.005), indicating reduced geometric distortion in the surrogate condition (Figure 5a).

**Figure 5:**
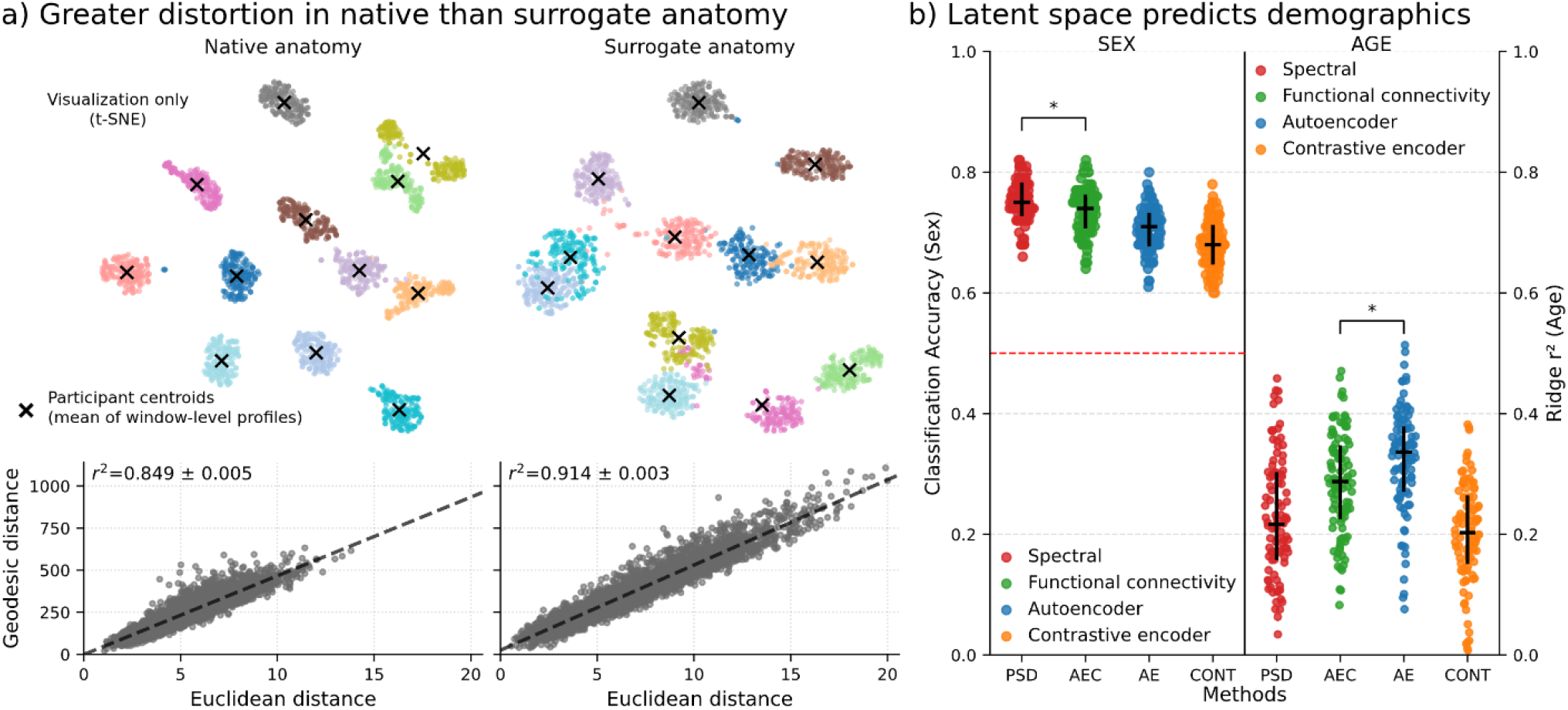
Latent space geometry and demographic associations. a) Greater geometric distortion in native than surrogate anatomy. Two-dimensional visualization of participant-level autoencoder profiles (visualization only; t-SNE) for native (left) and surrogate (right) anatomy in the Cam-CAN dataset. Each point represents a window-level embedding; crosses denote participant centroids (mean of window-level profiles). Bottom panels show the relationship between Euclidean and geodesic distances between participant centroids. Reduced deviation from linearity in the surrogate anatomy condition is reflected by a higher coefficient of determination (r^2^). b) Latent space predicts demographics. Sex classification accuracy (left) and age prediction performance (right; partial least-square regression r^2^) across profiling methods on 400 unseen Cam-CAN participants (100 random 75/25% train/test splits). Autoencoder-based profiles yield the most accurate age predictions, while spectral profiles perform best for sex classification. Dashed red line indicates chance-level performance for sex classification. Error bars denote quartiles across splits.

Consistent with this difference in geometry, age prediction using partial-least square regression was markedly more accurate in the surrogate anatomy space (r^2^ = 0.318 ± 0.082) than in the native anatomy space (r^2^ = 0.074 ± 0.071) for the autoencoder profiles. Sex classification accuracy was comparable across anatomical conditions (surrogate: 71.0 ± 4.0%; native: 70.5 ± 3.6%), with spectral profiles yielding the highest accuracy overall (73.5 ± 3.8%, p = 0.004). Autoencoder-based profiles achieved the most accurate age predictions among all methods (p = 0.002; Figure 5b).

## Discussion

Accumulating evidence indicates that resting-state brain activity expresses stable individual-specific structure^2,7–9,34^. Neurophysiological profiles derived from functional connectivity and spectral features can be heritable, relate to demographic or clinical variables, and be modulated by neurological disease^12–14^. However, reliance on model-free profiling constrains the space of candidate features and limits the ability to learn richer representations from large datasets.

Here, we introduced a participant-agnostic autoencoder framework for neurophysiological time series that extracts stable profiles directly from data. These profiles differentiate individuals using brief recordings, generalize above chance across recording sessions and remain robust when participant-specific anatomical information is withheld during source reconstruction.

### The Value of Unsupervised Profiling

Our approach differs from contrastive or other participant-supervised profiling models. By optimizing reconstruction fidelity rather than participant separability, the model is encouraged to learn the data distribution rather than a subspace shaped explicitly by labels or decision boundaries. This encourages the encoder to compress rich signal dynamics and may reduce reliance on features that are discriminative but biologically trivial. The risk of shortcut learning associated with participant-supervised objectives was illustrated by the contrastive baseline, which showed a substantially larger performance drop (86.9 percentage points) when generalizing across recording sessions than the other approaches (average drop, 46.9 percentage points). Reconstruction-based objectives can still encode confounds when those confounds reduce reconstruction error, but they are not explicitly incentivized to exploit them for participant separability.

This participant-agnostic framework also addresses a major barrier to scalable neurophysiological profiling: leveraging large, heterogeneous datasets without consistent participant labels. Model-free approaches are interpretable but rely on fixed mathematical transforms that do not improve with increasing cohort size. Supervised models can learn richer representations, but scaling them across datasets requires harmonized labels and careful control of site- and dataset-specific confounds. By removing the dependence on participant labels while retaining a learned transformation, our approach supports large-scale pretraining across heterogeneous cohorts and moves toward foundation-model strategies for neurophysiological profiling.

### Comparison to Previous Approaches

Autoencoder-based profiles were the most discriminative profiles of tested approaches in the Cam-CAN surrogate-anatomy setting and remained competitive in between-session profiling. Notably, the approach retained high performance when individual structural MRI data were unavailable and source reconstruction relied on a common anatomical template. Under this surrogate anatomy condition, profiling accuracy decreased by only 6.7 percentage points for the AE, compared with reductions of 12.6-19.8 percentage points for baseline methods. This robustness is relevant for applications where participant-specific anatomy is unavailable for practical, logistical or financial reasons.

Unlike model-free baselines, band-pass filtering did not preferentially enhance or degrade AE profiling accuracy for any low-frequency band below gamma. This relative invariance suggests that the autoencoder does not rely on a narrow spectral regime, but instead integrates information distributed across frequencies. Two non-exclusive mechanisms may underlie this behavior: redundancy of discriminative information across frequency bands, and sensitivity to broadband or aperiodic signal components that persist under band-pass filtering. This interpretation is consistent with recent findings that healthy adults can be differentiated by aperiodic components alone^35^. In contrast, rhythmic components may be more important for differentiating clinical populations, such as in Parkinson’s disease^14^, and developing cohorts^36^.

Support for this interpretation comes from the sensitivity analysis, which revealed structured spectral effects, and from the dimensionality reduction analysis, which showed negligible accuracy loss when retaining only approximately 5% of the latent dimensions. Models trained explicitly on narrowband signals might emphasize frequency-specific differences and could align more closely with prior band-limited profiling studies.

### Interpretability and Biological Features

The generative nature of the autoencoder enables direct interrogation of features associated with latent variability. Ordering latent components by participant-specific stability revealed a structured hierarchy, with leading components exhibiting high intraclass correlation. Combined with systematic occlusion, this structure allows latent dimensions to be mapped back to interpretable time-domain signals and derived feature spaces. Although our analyses focused on functional connectivity and spectral features to facilitate comparison with model-free baselines, the decoder-based approach is flexible: any feature derived from reconstructed time series, static or dynamic, can be interrogated without changing the training objective.

This capability is particularly relevant for transient and non-linear neural dynamics, such as bursting activity or neural avalanches, which have recently been implicated in individual variability and profiling^37,38^. Across the first five principal components, sensitivity analyses consistently highlighted dorsal attention networks in both connectivity and spectral domains. The topographic alignment with prior saliency findings based on ICC analyses in MEG^8,9^ is notable given the methodological differences between approaches. Although these effects are more spatially confined than those reported in recent large-scale cohort studies^36,39^, this likely reflects our focus on the first five principal components, which capture the dominant axes of participant variability.

### Demographic Associations and Translational Considerations

Autoencoder-based profiles predicted demographic variables with performance comparable to, or exceeding, model-free approaches, supporting the interpretation that they encode biologically meaningful information. Age prediction was notably more accurate using autoencoder profiles (r^2^ = 0.318). However, this performance depended on latent-space geometry: prediction in the native anatomy space showed increased inter-participant distortion and markedly reduced accuracy (r^2^ = 0.074). This result suggests that downstream regression tasks may require either explicit geometric constraints during training or selection of representations with intrinsically low distortion.

### Limitations and Future Directions

Relative to model-free baselines, the proposed architecture is more computationally and data intensive, and high-frequency activity was under-represented in reconstructed signals. Differentiation from brief recordings also raises the possibility that models capture stable technical artifacts in addition to neurophysiological traits. Although we demonstrate generalization across sessions, we did not test robustness across recording sites, devices or preprocessing pipelines, each of which can introduce stable dataset-specific structure. In particular, Cam-CAN and OMEGA differ in resting condition (eyes-closed vs. predominantly eyes open), which our design does not disentangle from session and site effects. Stable non-neural sources such as head motion and within-session head-position structure may also contribute to differentiation and were not explicitly controlled. The interpretability framework also remains indirect, relying on perturbation-based sensitivity rather than mechanistic explanations of latent variables.

Future work could explore model distillation or reduced-capacity architectures to lower computational demands. Pretraining on large multi-site datasets could improve robustness and enable generalization to unseen cohorts without retraining, consistent with emerging foundation-model paradigms^40–42^. Incorporating spectral or topographic priors may improve high-frequency and spatial fidelity, while decoder-based analyses could test how demographic or clinical variables modulate specific interpretable features. Ultimately, models trained to capture normative distributions of brain activity may support applications beyond differentiation, including quantifying disease-related deviations from typical neurophysiological structure^43,44^.

## Conclusion

We demonstrate that participant-agnostic unsupervised representation learning applied to cortical time series can extract stable, participant-specific neurophysiological profiles that generalize across sessions and anatomical constraints. By avoiding participant-supervised objectives and using a generative decoder for perturbation-based interpretability, the framework bridges interpretable model-free profiling and supervised representations that may be vulnerable to shortcut learning. Despite higher computational demands and reduced high-frequency fidelity, autoencoder-based profiles performed competitively with, and often exceeded, established baselines, supporting scalable and interpretable neurophysiological profiling.

## Methods

### Datasets and Demographics

We used data from the Cambridge Centre for Ageing and Neuroscience (Cam-CAN) repository^30^. Resting-state MEG recordings were acquired with eyes closed. After quality control (QC), the dataset comprised N = 599 participants, with approximately 9 min of data per participant. Mean age was 54.77 ± 18.31 years (range, 18.5-88.92 years). The cohort included 304 males and 295 females.

We additionally used data from the Open MEG Archives (OMEGA)^33^. Resting-state recordings were acquired with eyes open in most cases. After QC, the dataset comprised N = 324 participants, with approximately 5 min of data per participant. Mean age was 29.65 ± 12.27 years (range, 17.99-74.00 years). The cohort included 147 males and 177 females.

### MEG Preprocessing

Preprocessing pipelines differed between datasets but followed established MEG analysis practices^45,46^. Dataset-specific steps are detailed below.

#### Cam-CAN

Data were acquired at 1 kHz using a 306-channel Elekta Neuromag VectorView system (204 planar gradiometers, 102 magnetometers). Preprocessing was performed in Brainstorm^47^ (March 2021 release, MATLAB R2020b).

Line noise was removed using a 50 Hz notch-filter bank (10 harmonics). An additional 88 Hz notch filter was applied to remove a dataset-specific artifact. Signals were high-pass filtered at 0.3 Hz (FIR) to remove slow drifts. Cardiac, ocular and muscle artifacts were attenuated using signal-space projections (SSPs; 1-7 Hz for saccades, 40-400 Hz for muscle activity). Data were low-pass filtered at 75 Hz using a sixth-order Butterworth filter and downsampled to 150 Hz by FFT-based resampling.

Structural T1-weighted MRI volumes were segmented using FreeSurfer^48^ and coregistered to MEG sensor space using approximately 100 head points. Head models were computed using Brainstorm’s overlapping-spheres method. Source activity was estimated at 15,000 cortical dipoles using a linearly constrained minimum-variance (LCMV) beamformer with orientations constrained normal to the cortical surface. Source time series were parcellated to the Schaefer-200 atlas with Yeo-17 network labels^31,32^.

Signals within parcels were averaged with sign correction based on the dot product between source and parcel normals. Processed data were stored as single-precision tensors.

#### Native and Surrogate Anatomy Conditions

To assess the contribution of participant-specific anatomy to profiling performance, source reconstruction was performed under two conditions.

In the *native anatomy* condition, each participant’s own anatomy was used to compute both the LCMV spatial filter and the interpolation kernel to a common template.

In the surrogate anatomy condition, a single held-out participant’s anatomy was used to compute a fixed spatial filter and interpolation kernel that were applied to all participants. The surrogate anatomy was drawn at random from the cohort, and that participant was excluded from all training and evaluation sets. A single surrogate anatomy was used throughout. The analysis was not repeated with alternative surrogates. This manipulation does not remove all anatomical variability and should be interpreted as a robustness test rather than a complete removal of anatomical confounds.

Formally, let *X*_*p*_ ∈ ℝ^*MxT*^ denote the preprocessed sensor-space signals for participant *p*, with M sensors and T time points. For each participant, an LCMV spatial filter *W*_*p*_ ∈ ℝ^*SpxM*^ was computed in native source space, along with an interpolation kernel *I*_*p*_ ∈ ℝ^*STxSp*^ mapping to the ICBM152 template^49^. Here, *S*_*p*_ is the number of sources in participant *p*’s source space, and *S*_*T*_ is the number of sources in the template space.

Native anatomy condition: Source time series were obtained by applying *W*_*p*_ to *X*_*p*_ and interpolating with *I*_*p*_:

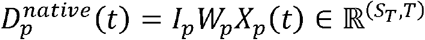

Surrogate anatomy condition: A fixed spatial filter *W*_*s*_ and interpolation kernel *I*_*s*_, derived from a held-out participant *s*, were applied to all *X*_*p*_:

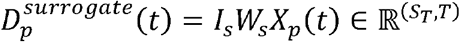

In both cases, source time series were subsequently parcellated to the Schaefer-200 atlas.

#### OMEGA

OMEGA data were acquired at 2.4 kHz using a 275-channel CTF axial-gradiometer system. Preprocessing was performed in MNE-Python^50^ (v1.10.0). Signals were band-pass filtered between 0.5 and 75 Hz (FIR), notched at 60, 120 and 180 Hz, and resampled to 150 Hz using FFT-based resampling. ECG and EOG artifacts were removed using SSPs.

MEG recordings were coregistered to the FreeSurfer fsaverage anatomy using participant-specific fiducials followed by iterative closest-point alignment. The forward model used MNE’s precomputed three-layer fsaverage BEM (inner skull, outer skull, and outer skin surfaces, each tessellated at ico-4 with 5120 vertices per surface). Sources were placed on a cortical surface source space constructed at ico-5 resolution, with 10,242 dipoles per hemisphere (approximate mean inter-source spacing of 3.1 mm). LCMV beamformers with unit-noise-gain normalization and orientation selection by maximal power were applied. Source time series were parcellated to the Schaefer-200 atlas and averaged with sign correction. Processed data were stored as single-precision tensors.

A subset of 78 OMEGA participants had multiple recording sessions. The mean inter-session interval was 115 days (5th-95th percentile, 9-300 days).

Because the two datasets were processed in different toolboxes, source-reconstruction details follow each toolbox’s established defaults and differ accordingly: Cam-CAN used an overlapping-spheres head model with LCMV orientations constrained normal to the cortical surface, whereas OMEGA used a three-layer BEM with unit-noise-gain LCMV normalization and orientation selected by maximal power. Because profiling was evaluated within each dataset separately, these cross-toolbox differences do not affect comparisons.

### Profiling

MEG time series were transformed into fixed-length feature vectors, or profiles, for participant differentiation.

For each participant *p*, two profiles 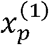 and 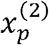 were constructed. An *N* × *N* similarity matrix was computed using Pearson correlation between all pairs *x*^(1)^ and *x*^(2)^.

Participant differentiation was considered correct when the maximum correlation in each row occurred on the diagonal. We report top-1 differentiation accuracy.

Within-session profiling: 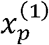 was computed from the first *L* seconds of a recording and 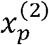 from the last *L* seconds.

Between-session profiling: 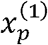 and 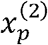 were computed from the first *L* seconds of two different sessions.

The effect of increasing *L* is investigated in the Effect of Segment Length on Differentiation Accuracy section. For the other experiments, *L* was fixed and shared across profiling approaches.

#### Model-Free Profiling

Functional connectivity profiles: AEC was computed by applying the Hilbert transform to each ROI time series and computing pairwise Pearson correlations between analytic envelopes. Profiles consisted of the flattened upper triangle of the matrix (19,000 dimensions).

Spectral profiles: PSD was estimated per ROI using Welch’s method^51^ (2-s windows, 50% overlap, Hanning window, nfft = 300, fs = 150 Hz), yielding a frequency resolution of *Δf* = *f*_*s*_/*n*_*fft*_ = 0.5 *Hz* (151 bins over 0-75 Hz). Profiles were formed by concatenating ROI spectra (200 ROIS × 151 bins = 30,200 dimensions).

#### Model-Based Profiling

Two encoders mapped input windows with *C* channels and *W* time points to *D* - dimensional latent vectors: an autoencoder (AE) and a contrastive encoder (CONT)^52^. Models were trained on non-overlapping 6-s windows (900 time points).

Profiles were extracted from multiple latent vectors *z*_*t*_. For segments longer than one window, we slid over the data, computed *z*_*t*_ and temporally pooled embeddings with step size *k* to suppress transient information:

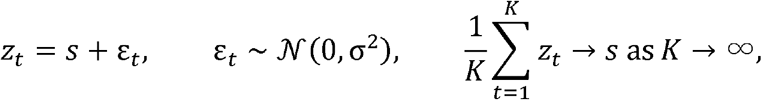

where *s* is the stable participant-specific signal, *ε*_*t*_ is a zero-mean noise term and *K* is the number of windows in a segment of length *L*.

Step sizes were set per method at the value beyond which differentiation accuracy saturated, as determined from the step-size analysis in Extended Data Figure 9: *k* = 1 sample for the AE and *k* = 150 samples (1 s) for CONT. Smaller step sizes increased computational cost without improving CONT accuracy, so the larger CONT step size reflects its saturation point.

### Model Architectures

#### Autoencoder

We adapted a KL-regularized variational autoencoder inspired by latent-diffusion VAEs^53^ to spatiotemporal MEG windows (Figure 6). Inputs were tensors of shape 1 × *ROIs* × *time*. The encoder comprised stacked 2-D convolutional blocks with Group Normalization^54^ (32 groups), SiLU activations^55^ and dropout (p = 0.1). Single-head self-attention modules^56^ were inserted at selected resolutions.

**Figure 6:**
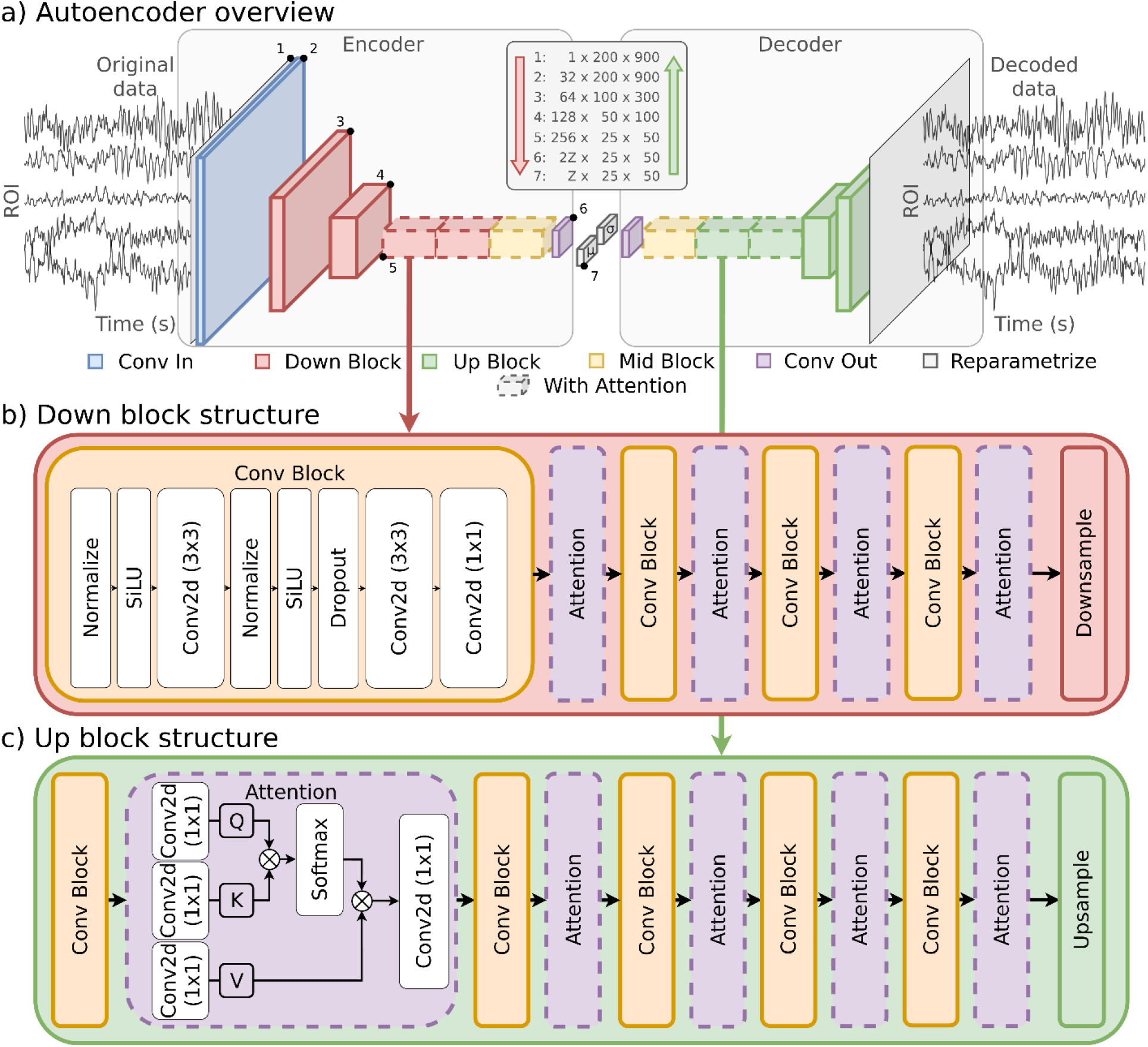
Autoencoder architecture used for representation learning. a) Autoencoder overview. Convolutional encoder-decoder architecture applied to source-space MEG time-series windows (200 ROIs × 900 time points). The encoder progressively reduces spatial and temporal resolution using convolutional down blocks, producing a latent representation sampled via reparameterization and symmetrically decoded to reconstruct the input. Attention modules are inserted at lower-resolution stages to improve modeling of long-range dependencies. b) <u>Down block structure</u>. Each down block comprises convolutional layers with group normalization, SiLU activation, and dropout, optionally followed by self-attention, before downsampling. c) Up block structure. Each up block mirrors the down block, with upsampling followed by convolutional layers and optional self-attention. Attention uses a standard query-key-value formulation with softmax normalization. Architectural components are generic and were not tuned for participant differentiation or demographic prediction.

Downsampling used anisotropic strided convolutions; upsampling used linear interpolation followed by convolution. The encoder parameterized a diagonal Gaussian posterior sampled with the reparameterization trick^57,58^. A symmetric decoder mirrored the encoder. Training optimized a standard VAE objective^59^ (MSE plus -weighted KL divergence).

The pooled latent dimensionality used for profiling was . The AE contained approximately 37 million trainable parameters.

#### Contrastive Encoder

The contrastive encoder shared the AE encoder backbone. The VAE head was replaced by a 1 × 1 projection (stride 2) that reduced channel dimensionality from 2*z*_*ch*_ to *z*_*ch*_. Training used a Siamese architecture with cosine embedding loss (margin *m* =0.5) for window pairs. Positive pairs () comprised two windows from the same participant; negative pairs (*y* = + 1) comprised windows from different participants.

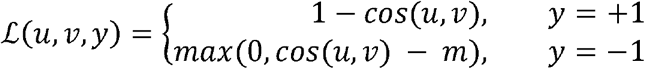

To confirm that the pairwise objective was not limited by convergence instability associated with contrastive losses^60,61^, we also tested a supervised classification baseline. The encoder output was projected through a linear layer to classes corresponding to the training participants and optimized with cross-entropy loss. For inference on unseen participants, the classification head was discarded and the encoder output served as the profile. Both formulations yielded comparable differentiation accuracy (Extended Data Figure 9); main-text results therefore report the cosine embedding loss baseline.

Output dimensionality was 20,000, with approximately 16 million trainable parameters.

#### Fine-Tuned Autoencoder

After pretraining, the AE was fine-tuned for 10 epochs (approximately 10% of pretraining duration) on unlabeled evaluation data using the same reconstruction objective. Participant labels and demographic variables were not used during fine-tuning nor were they used during pre-training. This fine-tuning extends the reconstruction objective to the unlabeled target recordings to adapt the latent space to the target dataset’s statistics; it does not introduce identity information. However, because fine-tuning exposes the AE to the full evaluation recordings under a reconstruction objective, the fine-tuned AE’s performance at segment lengths less than the full length partially reflects familiarity with the underlying recordings. The pre-trained AE is therefore the appropriate point of comparison for data-efficiency claims at short segment lengths. The fine-tuned AE is reported as an upper-bound condition. No fine-tuning was applied to CONT as it requires an identity signal during training.

#### Model Training

For each dataset, 200 participants were assigned to the training set. For OMEGA, these were drawn from single-session participants, reserving multi-session participants for between-session testing. A 5% validation subset was randomly drawn from the training set and used to monitor convergence. Random seeds were drawn once and saved. The autoencoder used seed 5885 and the contrastive encoder used seed 2692. The number of epochs was fixed past the empirically observed point of convergence. Both models were trained for sufficient time to converge, and the converged checkpoint was used for all evaluations.

All models were implemented in PyTorch^62^ v2.7.1 and trained on an RTX 4080 SUPER GPU (16 GB VRAM). Gradient checkpointing was used for all architectures to reduce memory usage. Full training of each autoencoder took approximately 15 h; contrastive encoders required approximately 11 h.

Networks were optimized using AdamW^63^ with a fixed learning rate of 3 × 10^−4^ and default parameters (*β*_1_ = 0.9, *β*_2_ = 0.999). We used a batch size of 4 for all training runs. Detailed hyperparameter configurations for each dataset and model variant are provided in Extended Data Table 1.

### Experimental Setup

#### Autoencoder Output Validation

We first evaluated whether the autoencoder preserved key properties of the underlying MEG signals. For each participant in the Cam-CAN test set (N = 100), 10 non-overlapping windows of 6 s were encoded and decoded.

Time series fidelity: All decoded windows were concatenated per participant, channels were flattened, and the Pearson correlation between original and decoded time series was computed. We report mean ± SD across participants.

Functional connectivity fidelity: AEC, as defined in Model-free profiling, was computed on original and decoded windows. Pearson correlation between the upper triangles of the AEC matrices was computed per window and averaged per participant. We report mean ± SD across participants.

Peak alpha frequency (PAF): PSD was averaged across 10 consecutive windows (60 s). The PAF was defined as the discrete frequency bin with the maximum power within the 8-13 Hz band, extracted from the raw spectrum for each ROI. Pearson correlation between original and decoded PAF topographies was computed per participant. We report mean ± SD across participants.

Spectral fidelity: PSD, as defined in Model-free profiling, was estimated per ROI and grouped by Yeo-17 networks. To quantify evidence for similarity between original and decoded power spectra, we computed Bayes factors (BF01) using Bayesian paired t-tests at each frequency bin. We used *BF*_01_ ≥ 3 as moderate evidence in favor of the null hypothesis^64,65^.

Time-frequency structure: Time-frequency decompositions were computed using Morlet wavelets (1–30 Hz, 3 cycles) and used for qualitative comparison of temporal–spectral structure.

### Profiling Experiments

We trained one autoencoder and one contrastive encoder per dataset. The AE was additionally fine-tuned on the test cohort (fine-tuned AE).

Cam-CAN: AE, fine-tuned AE and CONT were trained in the native anatomy space. Models were evaluated on a single independent test set of 100 participants reconstructed in both native and surrogate anatomy space. The two conditions therefore comprise the same participants reconstructed two ways.

OMEGA: AE and CONT were trained on within-session data. The fine-tuned AE was further optimized using unlabeled data from multiple full sessions with reconstruction loss only. Models were evaluated on two test sets of 78 participants: one for within-session profiling and one for between-session profiling.

Unless otherwise specified, reported values correspond to mean ± SD over 100 repetitions of 90% participant subsampling without replacement.

#### Effect of Segment Length on Differentiation Accuracy

Profiles were computed using segment lengths *L* ∈ [6,120] s in 1-s increments. For each L, differentiation accuracy was computed for AEC, PSD, AE, fine-tuned AE and CONT. Cam-CAN results correspond to the surrogate anatomy condition; OMEGA results correspond to between-session profiling.

For each combination of method and segment length, differentiation accuracy was estimated from 100 bootstrap iterations, each formed by sampling 90% of subjects without replacement. Bootstrap resamples were drawn independently for each method. Between-method differences at specific segment lengths were tested with a one-sided Welch’s two-sample t-test. A one-sided Welch’s test was used because the alternative hypothesis was directional and specified a priori: the autoencoder was predicted to exceed the baseline methods at sufficiently long segments, which is the hypothesis motivating the segment-length analysis. To control for the multiple tests across the 115 segment lengths within each pairwise method comparison, raw p-values were Bonferroni-adjusted by multiplication by the number of segment lengths, and the *α* =0.05 significance threshold was used.

#### Robustness to Anatomy Alignment

Models trained on Cam-CAN data reconstructed in the native anatomy condition were evaluated on data reconstructed in the surrogate anatomy condition. Segment length was fixed at *L*= 120 s.

#### Generalizability to Between-Session Profiling

Models trained on OMEGA within-session data were evaluated on between-session data using *L* =120 s segments.

#### Effect of Post Hoc Latent Dimensionality Reduction

From the 20,000-dimensional autoencoder latent representation, we randomly subsampled *d* ∈ {1024,2048,4096,8192,16384} dimensions (100 draws per d) and recomputed differentiation accuracy. This procedure was repeated for both datasets and all evaluation conditions using *L*= 120 s segments.

#### Frequency Band Contributions

Before profiling, input signals were band-pass filtered into canonical frequency bands: broadband (0.5-45 Hz), delta (0.5-4 Hz), theta (4-8 Hz), alpha (8-13 Hz), beta (13-30 Hz) and gamma (30-45 Hz). Models were not retrained for individual bands.

Cam-CAN results correspond to the *surrogate anatomy* condition. OMEGA results correspond to *between-session* profiling. Segment length was *L* = 120s.

### Interpretability Experiments

Interpretability of latent representations is essential for neurophysiological profiling. Encoder-only models lack a generative inverse; the autoencoder decoder enables perturbation-based sensitivity analyses in interpretable feature spaces.

#### Latent Space Stability Analysis

PCA^66^ was fitted to all training latent vectors (190 participants, approximately 30 windows per participant). Components explaining 95% of the variance were retained. Test latent vectors were projected into the same PCA basis.

For each principal component, *ICC* (3,1) ^67,68^ was computed across windows grouped by participant to quantify participant-specific stability. A least-squares fit of the form *ICC* (*k*) =*Ck* ^-α^ was used to model stability as a function of component index.

For each participant, two windows *a* and *b* were encoded into the PCA basis (*s*_*a*_, *s*_*b*_). For component *k*, the value *s*_*a*_ [*k*] was set to zero, decoded and re-encoded. Errors relative to the zeroed target, original latent and same-participant latent were computed as:

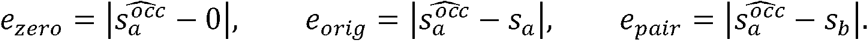

A non-scrambling decoder is characterized by low error relative to the zeroed target and larger errors relative to the other two comparisons^69^.

#### Decoder-based Feature Sensitivity Analysis

A test set of N = 397 participants yielded 5,226 6-s windows. Each window was encoded, projected into the training PCA basis, decoded and mapped to interpretable feature spaces. We analyzed AEC (200 × 200 features) and PSD (200 × 151 features). No features with zero participant-specific standard deviation were encountered. The pipeline raises an error on such cases.

For each participant and feature *f*, the base distribution consisted of feature values across that participant’s windows, with standard deviation *sd*_*f*_ . For window *a, x*_*f*_ (*a*) denotes the base feature value. After zeroing component *k*, decoding and feature extraction, the occluded value 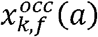 was obtained. Feature sensitivity was computed as the normalized deviation relative to the base distribution, with per-window feature sensitivity defined as:

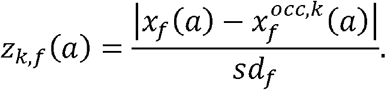

Feature sensitivities were averaged across all windows and participants, yielding one sensitivity value per AEC entry and per PSD bin.

#### Demographic Associations

Profiles were computed per participant as the mean of 10 non-overlapping 6-s windows. For visualization, t-SNE^70^ was applied to AE profiles (2-D, perplexity = 30).

To assess latent-space geometry, participant centroids were computed, pairwise Euclidean distances were calculated, and geodesic distances were estimated as shortest paths on a *k*-nearest-neighbor graph (*k* = 10, fixed a priori; edge weights = Euclidean). A least-squares model *d*_*geo*_ = *β*_0_ + *β*_1_ *d*_*Euc*_ was fitted, and the coefficient of determination (r^2^) was reported. Higher r^2^ indicates a more linear relationship between Euclidean and geodesic distances.

For demographic prediction, N = 400 unseen Cam-CAN participants were used. We performed 100 random 75/25% train/test splits, with features standardized using training-set means and standard deviations.

Age was predicted using partial least-squares regression with two components and accuracy was quantified using r^2^. Sex was predicted using L2-penalized logistic regression and quantified using top-1 classification accuracy.

Significance was assessed by comparing the best-performing method to its runner up with a two-sided Welch’s two-sample t-test. To control for the two demographic variables tested, reported p-values are Bonferroni-adjusted by a factor of two.

For each profiling method (AEC, PSD, AE, CONT), we report the distribution of performance metrics across the 100 splits.

## Data Availability

Raw data are available from the Cam-CAN^30^ and OMEGA^33^ repositories under their respective access policies (Cam-CAN: https://opendata.mrc-cbu.cam.ac.uk/projects/camcan/; OMEGA: https://www.mcgill.ca/bic/omega-registration).

Processed data generated are available from the corresponding author upon reasonable request.

Trained model weights are not redistributed because they derive from data governed by the Cam-CAN and OMEGA access agreements. Both datasets are available under those agreements and all weights can be regenerated from the released code.

## Code Availability

All analysis and modeling code is available at https://github.com/mLapatrie/aefp. (Latest commit: 13bf9f285671521476d3ef9863421c3bfd9b5e6d).

## Acknowledgments

S.B. was supported by a Discovery grant from the Natural Sciences and Engineering Research Council of Canada (436355-13), the CIHR Canada Research Chair in Neural Dynamics of Brain Systems (CRC-2017-00311), the NIH (R01-EB026299-05), and the Canadian Institutes of Health Research (202503PJT-542788). M.L. was supported by a Tanenbaum Open Science Institute Research Award. J.d.S.C. was supported by a doctoral fellowship from NSERC. We thank Doris Hua for assistance with the visual layout of Figure 1, and Elisabeth Jasmin for editorial assistance with the manuscript.

## Author Contributions

M.L., J.d.S.C., I.A., and S.B. conceptualized and designed the study. M.L. and J.d.S.C. curated and processed the data. M.L. developed the software and performed the formal analysis. I.A. validated the analysis code. M.L. created the visualizations. M.L. wrote the original draft of the manuscript. All authors contributed to the interpretation of the results and reviewed and edited the manuscript. S.B. supervised the project.

## Extended Data

**Extended Data Table 1:** Hyperparameters of models. Configuration details for the Autoencoder (AE) and Contrastive Encoder models used for the Cam-CAN and OMEGA datasets. Fine-tuned AE models used the same architecture and training parameters as the base AE but were additionally optimized on the relevant unlabeled evaluation subsets.

| Parameter | Cam-CAN |  | OMEGA |  |
| --- | --- | --- | --- | --- |
|  | AE | Contrastive | AE | Contrastive |
| *Training* |  |  |  |  |
| Batch Size | 4 | 4 | 4 | 4 |
| Learning Rate | $3 \times 10^{-4}$ | $3 \times 10^{-4}$ | $3 \times 10^{-4}$ | $3 \times 10^{-4}$ |
| Epochs | 100 | 100 | 100 | 100 |
| Optimizer | AdamW | AdamW | AdamW | AdamW |
| Loss Function | MSE + KL | Cosine Embedding | MSE + KL | Cosine Embedding |
| KL Weight (max) | $1 \times 10^{-5}$ | - | $1 \times 10^{-6}$ | - |
| Contrastive Margin | - | 0.5 | - | 0.5 |
| *Architecture* |  |  |  |  |
| Input Shape | 200 x 900 | 200 x 900 | 200 x 900 | 200 x 900 |
| Base Channels | 32 | 32 | 32 | 32 |
| Channel Multipliers | [2, 4, 8, 8] | [2, 4, 8, 8] | [2, 4, 8, 8] | [2, 4, 8, 8] |
| Latent Channels | 16 | 16 | 16 | 16 |
| Res. Blocks per Level | 4 | 4 | 4 | 4 |
| Attention Levels | [2, 3, 4, 5] | [2, 3, 4, 5] | [2, 3, 4, 5] | [2, 3, 4, 5] |
| Dropout | 0.1 | 0.1 | 0.1 | 0.1 |
| Strides | $\begin{bmatrix} 2,3 \\ 2,3 \\ 2,3 \\ 2,3 \end{bmatrix}$ | $\begin{bmatrix} 2,3 \\ 2,3 \\ 2,3 \\ 2,3 \end{bmatrix}$ | $\begin{bmatrix} 2,3 \\ 2,3 \\ 2,3 \\ 2,3 \end{bmatrix}$ | $\begin{bmatrix} 2,3 \\ 2,3 \\ 2,3 \\ 2,3 \end{bmatrix}$ |

**Extended Data Figure 1:**
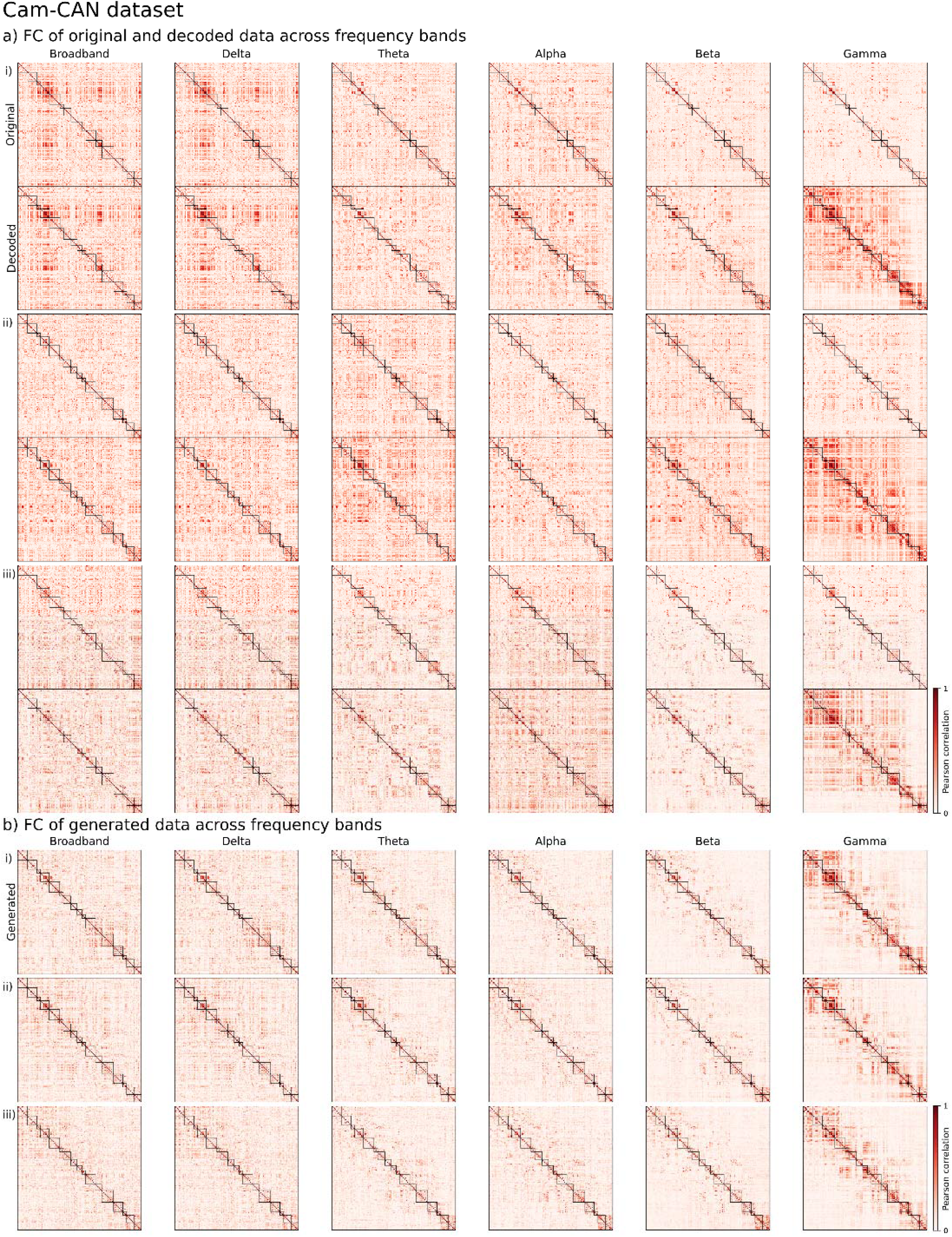
Functional connectivity structure across frequency bands in the Cam-CAN dataset. **a) Original and decoded data**. AEC matrices computed from source-level MEG signals across frequency bands (broadband, delta, theta, alpha, beta, gamma) for three representative participants (i-iii). For each participant, the original AEC matrix (top) is shown with the corresponding matrix computed from decoded signals (bottom). Values represent Pearson correlations between regional envelopes. Color scales are band-specific and shared between original and decoded matrices within each band. **b) Generated data**. AEC matrices computed from signals generated by the decoder after sampling from the learned latent distribution, shown for the same frequency bands and representative participants. Generated connectivity patterns exhibit structured organization similar to original and decoded data. Gamma-band results are shown for completeness and were not used in downstream profiling analyses.

**Extended Data Figure 2:**
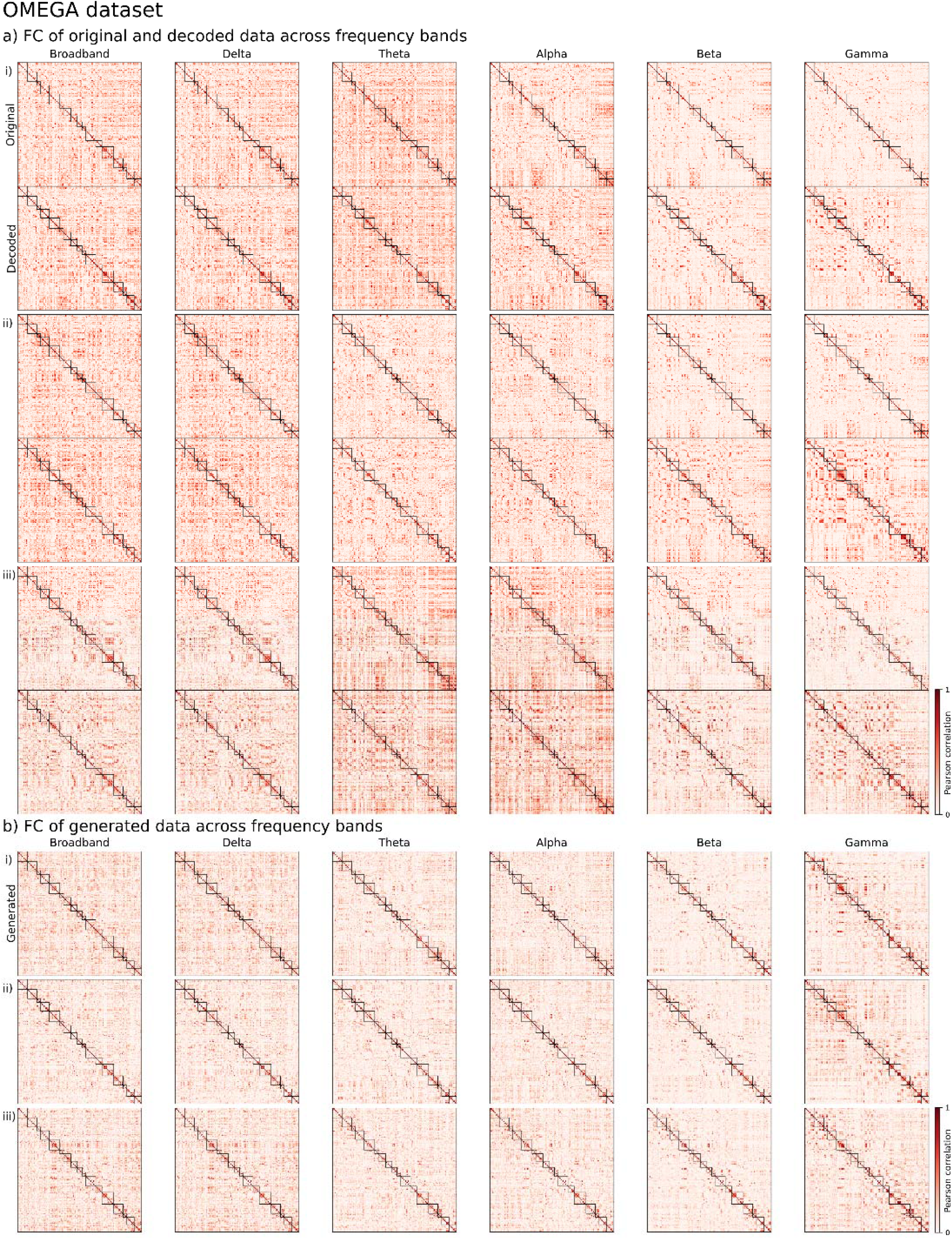
Functional connectivity structure across frequency bands in the OMEGA dataset. **a) Original and decoded data**. AEC matrices computed from source-level MEG signals across frequency bands (broadband, delta, theta, alpha, beta, gamma) for three representative OMEGA participants (i-iii). For each participant, original AEC matrices (top) are shown with matrices computed from decoded signals (bottom). Values represent Pearson correlations between regional envelopes. Color scales are band-specific and shared between original and decoded matrices within each band. **b) Generated data**. AEC matrices computed from signals generated by the decoder after sampling from the learned latent distribution, shown for the same frequency bands and representative participants. Generated connectivity patterns exhibit structured organization similar to original and decoded data. Gamma-band results are shown for completeness and were not used in downstream profiling analyses.

**Extended Data Figure 3:**
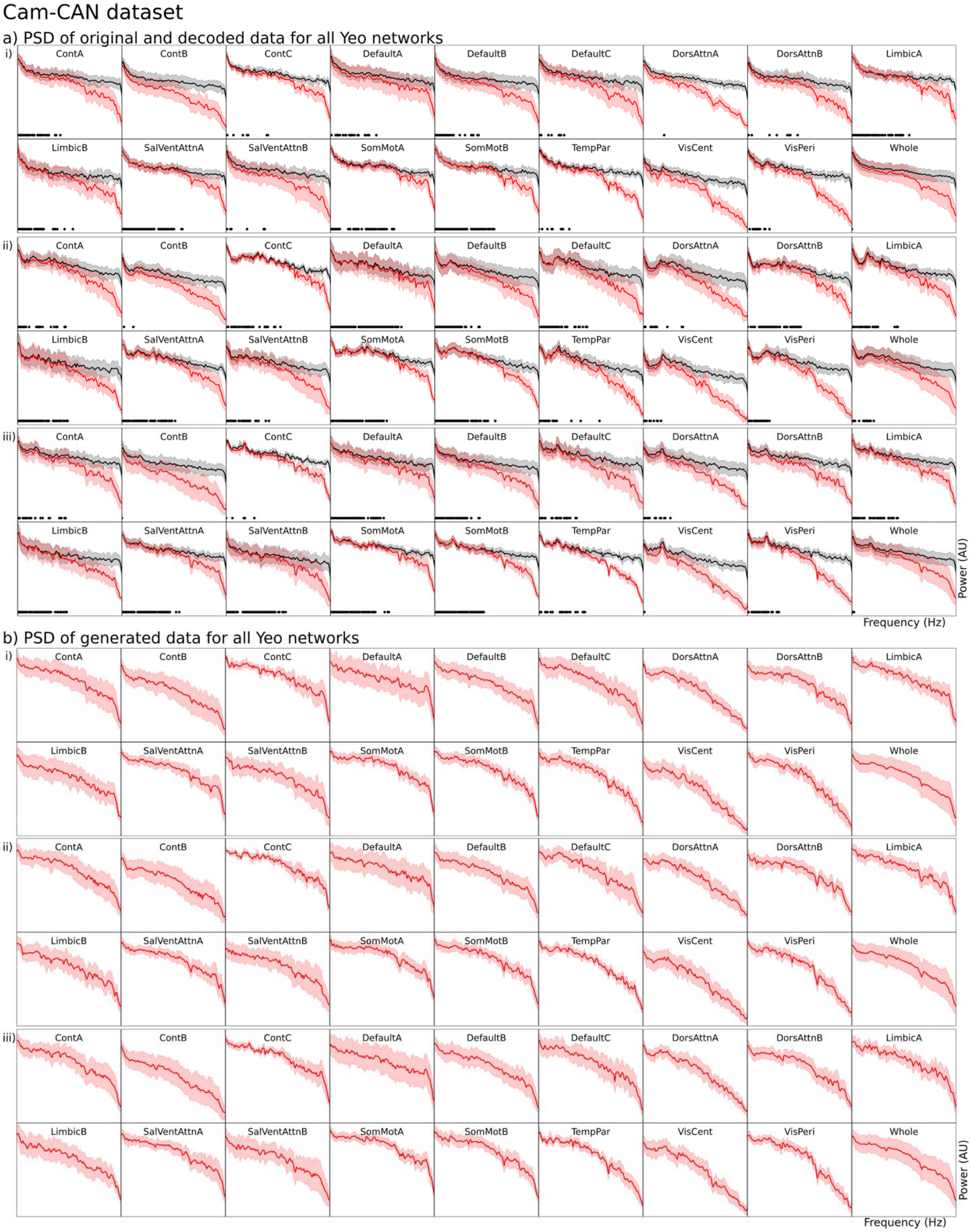
Power spectral density across Yeo-17 networks in the Cam-CAN dataset. a) Original and decoded data. PSD estimates for source-level MEG signals, shown for all Yeo-17 networks and three representative participants (i-iii). Black curves denote original data; red curves denote decoded signals. Shaded regions indicate variability across ROIs within each network. Horizontal black markers indicate frequencies with substantial evidence of similarity between original and decoded PSDs (BF01 >= 3). b) Generated data. PSDs computed from signals generated by the decoder after sampling from the learned latent distribution, shown for the same networks and representative participants. Generated spectra exhibit realistic broadband structure and network-specific profiles. Generated data are shown for qualitative assessment and were not used in profiling analyses.

**Extended Data Figure 4:**
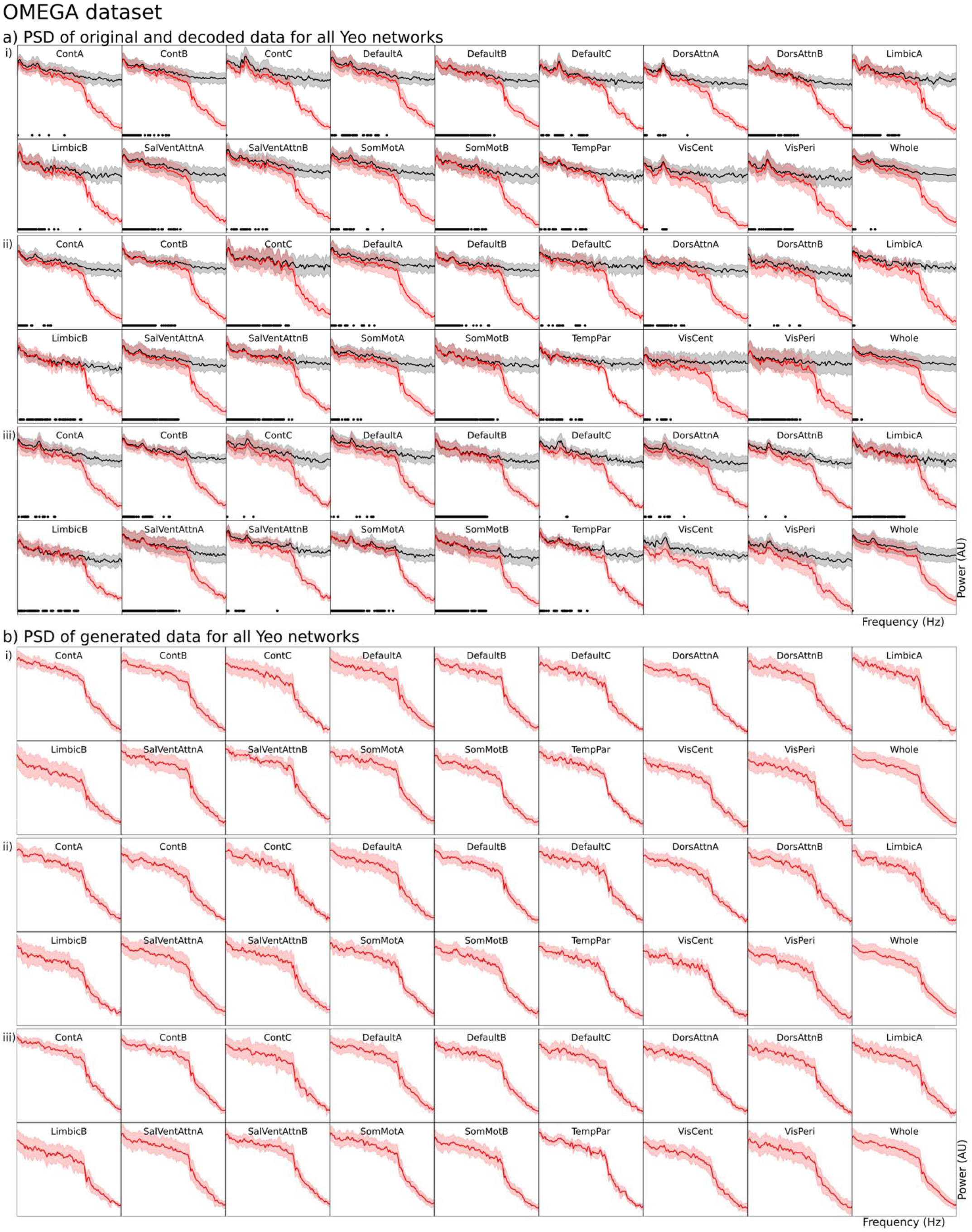
Power spectral density across Yeo-17 networks in the OMEGA dataset. a) Original and decoded data. PSD estimates for source-level MEG signals across all Yeo-17 networks and three representative OMEGA participants (i-iii). Black curves denote original data; red curves denote decoded signals. Shaded regions indicate variability across ROIs within each network. Horizontal black markers indicate frequencies with substantial evidence of similarity between original and decoded PSDs (BF01 >= 3). b) Generated data. PSDs computed from signals generated by the decoder after sampling from the learned latent distribution, shown for the same networks and representative participants. Generated spectra exhibit realistic broadband structure and network-specific profiles. Generated data are shown for qualitative assessment only and were not used in profiling analyses.

**Extended Data Figure 5:**
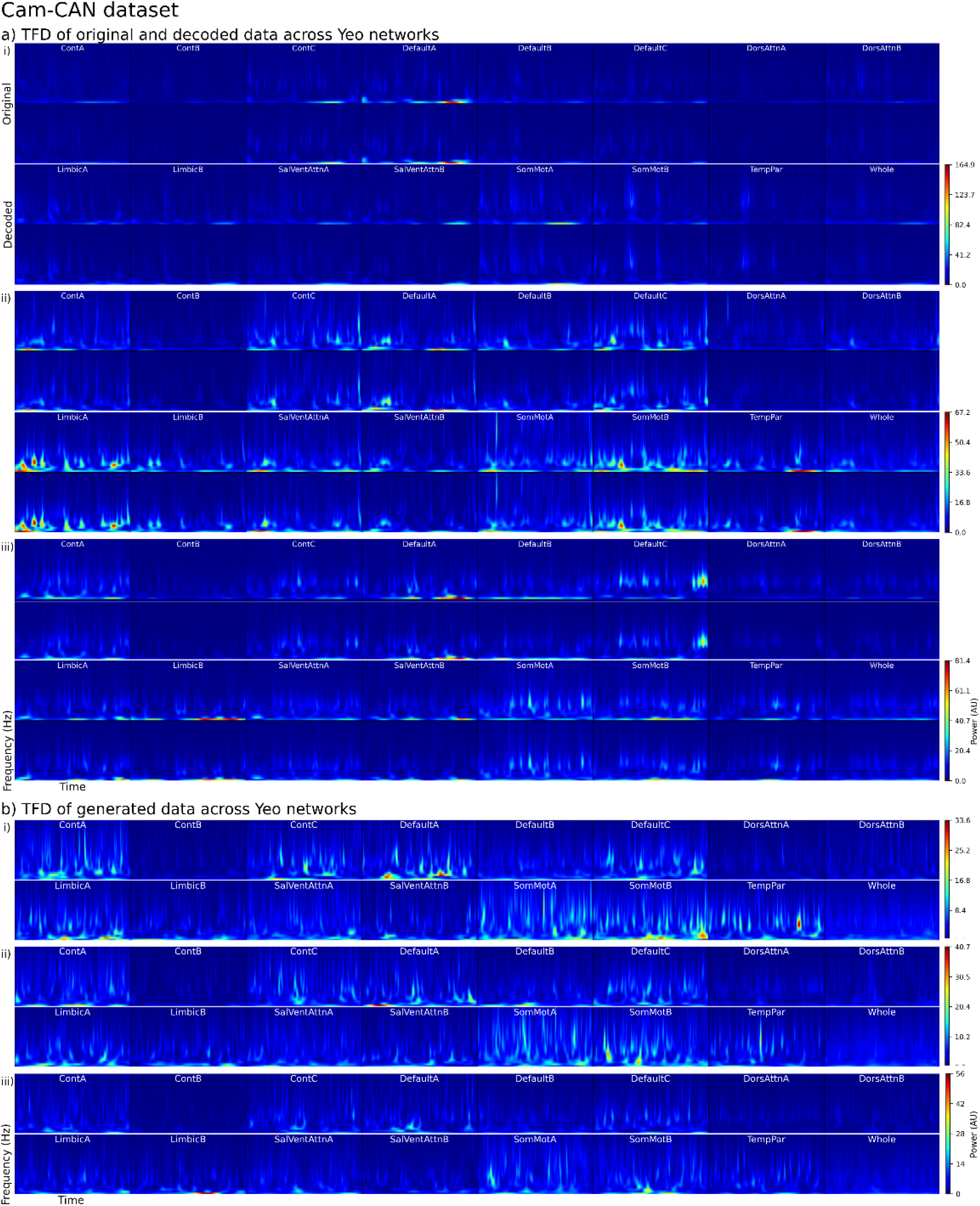
Time–frequency structure across Yeo-17 networks. **a) Original and decoded data**. Time-frequency decompositions (Morlet wavelets, 1-30 Hz, 3 cycles) of source-level MEG signals, shown for all Yeo-17 networks and three representative participants (i-iii). For each network, original data (top) are shown with decoded signals (bottom). Time-frequency representations are averaged across ROIs within each network. Color scales are normalized within each panel. Panels are shown for qualitative comparison only. **b) Generated data**. Time-frequency decompositions computed from signals generated by the decoder after sampling from the learned latent distribution, shown for the same networks and representative participants. Generated signals exhibit structured temporal-spectral patterns similar to those observed in original and decoded data. Generated data are shown for qualitative assessment only and were not used in downstream profiling analyses.

**Extended Data Figure 6:**
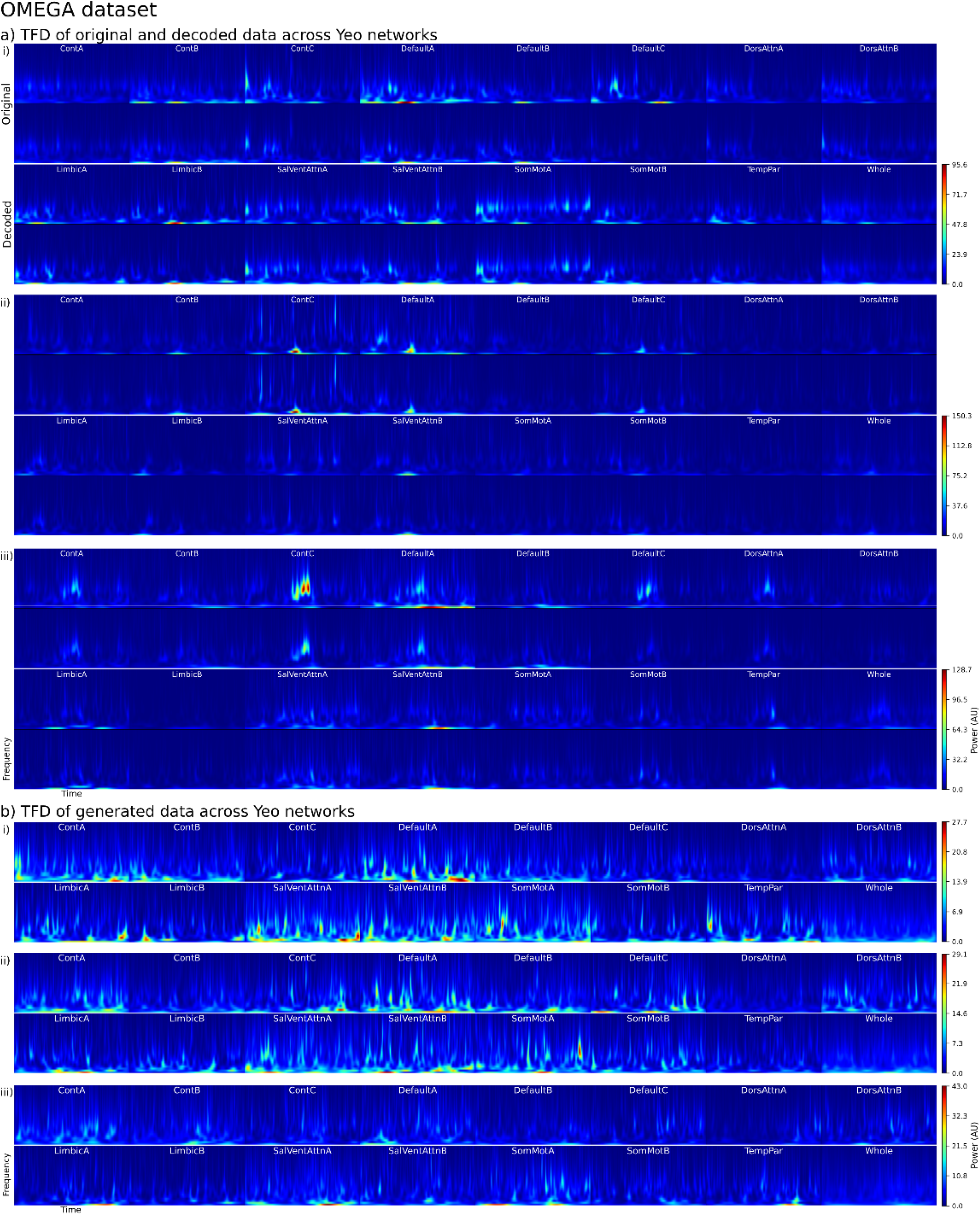
Time–frequency structure across Yeo-17 networks in the OMEGA dataset. **a) Original and decoded data**. Time-frequency decompositions (Morlet wavelets, 1-30 Hz, 3 cycles) of source-level MEG signals, shown for all Yeo-17 networks and three representative OMEGA participants (i-iii). For each network, original data (top) are shown with decoded signals (bottom). Time-frequency representations are averaged across ROIs within each network. Color scales are normalized within each panel. Panels are shown for qualitative comparison only. **b) Generated data**. Time-frequency decompositions computed from signals generated by the decoder after sampling from the learned latent distribution, shown for the same networks and representative participants. Generated signals exhibit structured temporal-spectral patterns similar to those observed in original and decoded data. Generated data are shown for qualitative assessment only and were not used in profiling analyses.

**Extended Data Figure 7:**
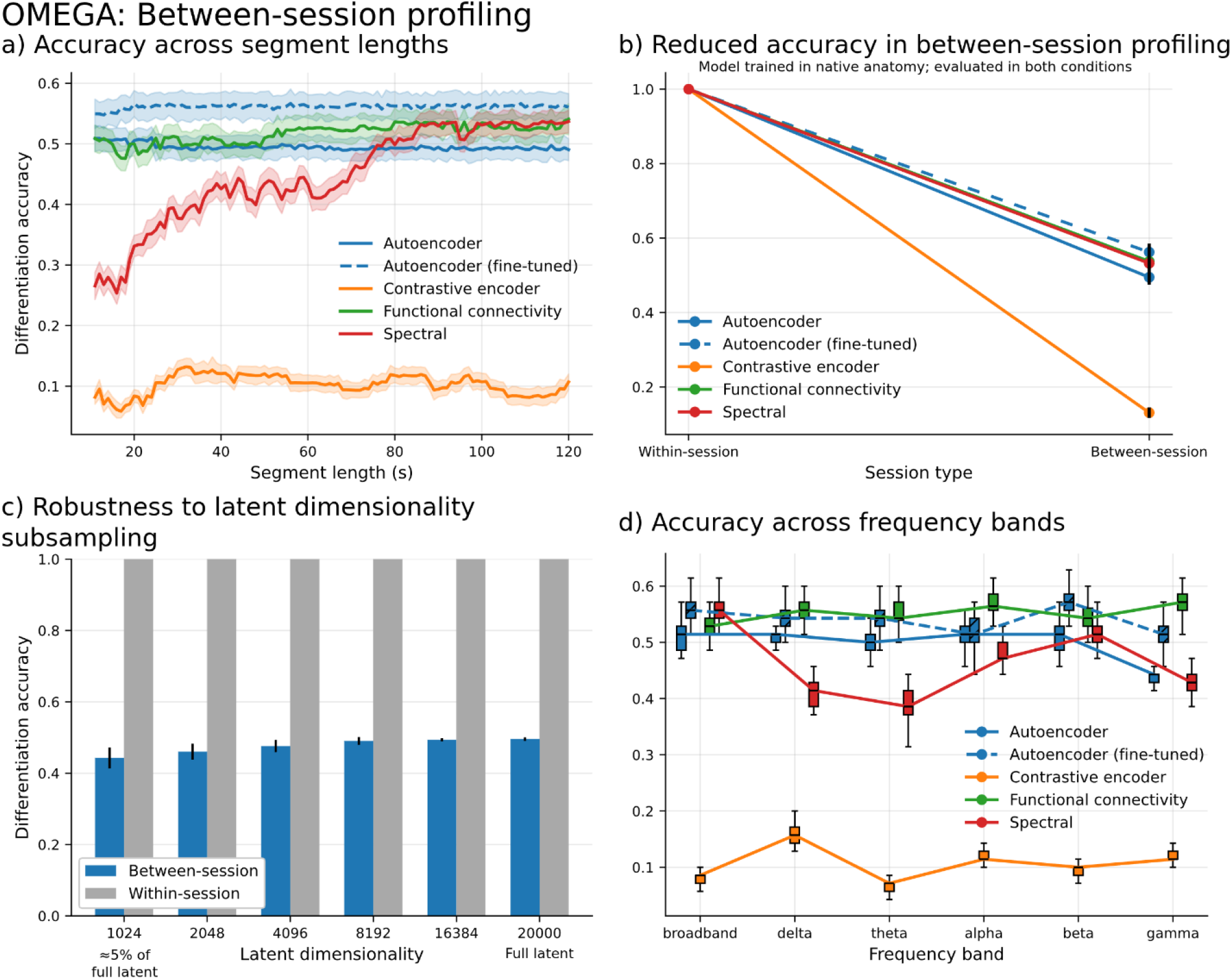
Between-session participant differentiation in the OMEGA dataset. **a)** Accuracy across segment length. Top-1 participant differentiation accuracy as a function of recording duration (10-120 s), computed by correlating profiles derived from different recording sessions of the same participants (N = 78). Curves show mean accuracy; shaded bands indicate bootstrap SD over 100 resamples of 90% of participants. Absolute accuracy is reduced relative to within-session profiling due to session variability, but relative method ordering is preserved. **b)** Reduced accuracy under between-session profiling. Differentiation accuracy for within-session and between-session profiling using 120-s segments. All models were trained on within-session data only, except the fine-tuned AE, which was adapted with reconstruction loss only. Between-session accuracy decreases for all methods, with contrastive learning showing the largest degradation. **c)** Robustness to latent dimensionality subsampling. Differentiation accuracy after randomly subsampling the autoencoder latent space at multiple dimensionalities (100 draws per dimensionality). Compression robustness persists under between-session conditions. **d)** Accuracy across frequency bands. Differentiation accuracy at 120 s after band-pass filtering inputs into canonical frequency bands. Autoencoder-based profiles remain relatively invariant across bands. Gamma-band results are shown for completeness.

**Extended Data Figure 8:**
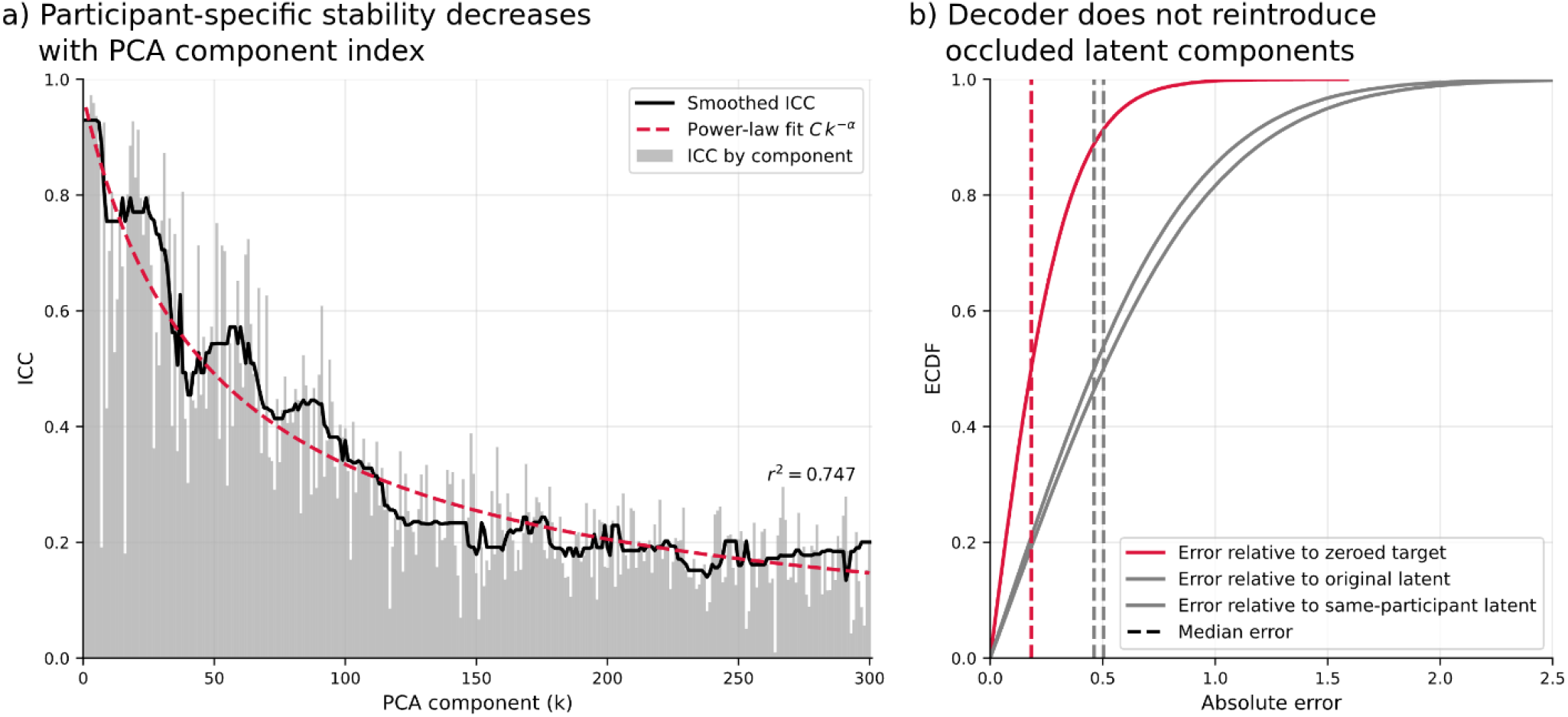
Stability structure of the autoencoder latent space. **a) Participant-specific stability decreases with PCA component index**. Intraclass correlation coefficients (ICC(3,1)) computed for each principal component of the autoencoder latent space across windows grouped by participant. Grey bars indicate ICC values per component; the black curve shows a smoothed trend. A power-law fit (ICC(k) = Ck^−α^) captures the decay of stability with component index (r^2^ = 0.747), indicating that leading components encode more participant-specific structure. **b) Decoder does not reintroduce occluded latent components**. Empirical cumulative distribution functions (ECDFs) of absolute reconstruction error following occlusion of individual principal components. Errors are shown relative to the zeroed target (red), the original latent value (light grey), and another latent vector from the same participant (dark grey). Vertical dashed lines denote median absolute error. Re-encoded representations remain closer to the zeroed target than to the original or same-participant latents, indicating that the decoder does not reconstruct occluded components.

**Extended Data Figure 9:**
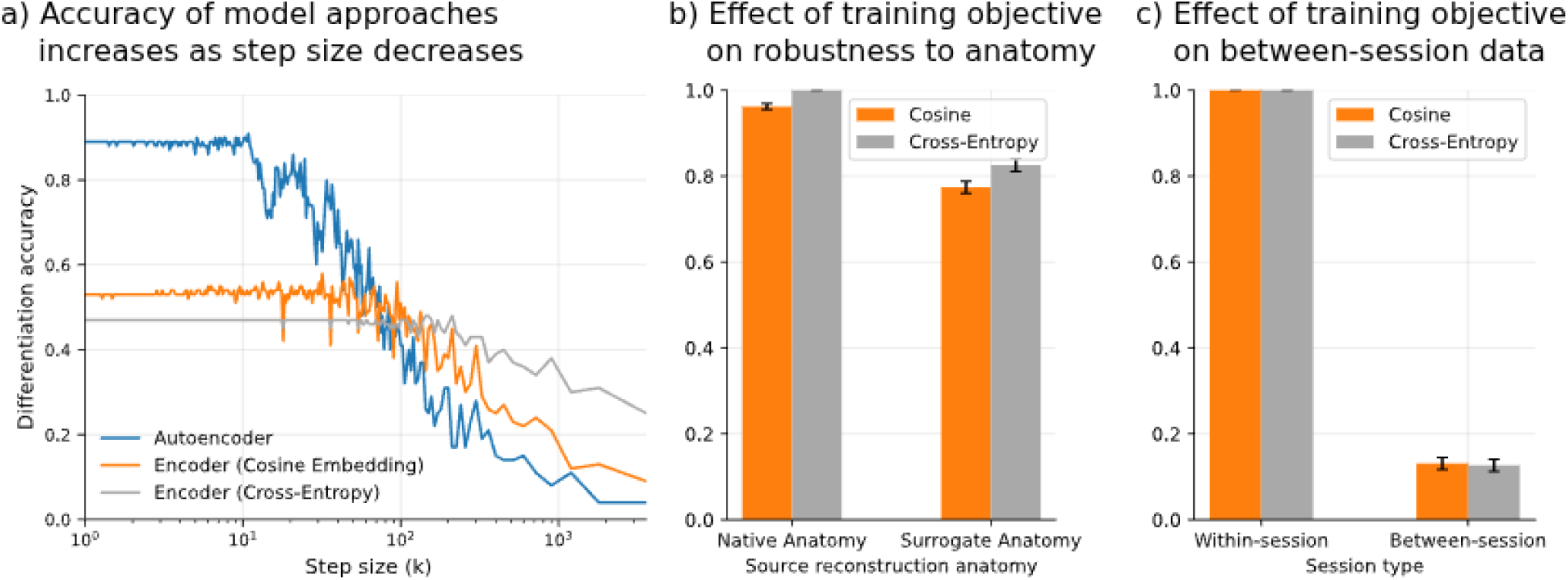
Choice of model-based profiling parameters. **a) Accuracy of model-based approaches increases as step size decreases**. Differentiation accuracy of model-based approaches as a function of the step size used for temporal pooling of latent embeddings. Models were evaluated on 30-s segments from the Cam-CAN surrogate anatomy condition. Smaller step sizes increase computational cost but yield higher accuracy by better estimating the stable participant-specific signal. Autoencoder accuracy mostly saturates below step sizes of 100 time points (approximately 0.67 s); encoder accuracy (cosine embedding and cross-entropy losses) saturates below step sizes of 200 time points (approximately 1.33 s). **b) Effect of training objective on robustness to anatomy alignment**. Native and surrogate anatomy differentiation accuracy on 120-s segments for models trained on Cam-CAN native-anatomy data. Encoders were trained using either cosine embedding loss or cross-entropy loss. Both training objectives yield similar robustness to anatomy mismatch. **c) Effect of training objective on between-session profiling**. Within- and between-session differentiation accuracy on 120-s segments for models trained on OMEGA within-session data using either cosine embedding loss or cross-entropy loss. Both training objectives yield similar drops in accuracy when evaluated between sessions.

